# Progressive overfilling of readily releasable pool underlies short-term facilitation at recurrent excitatory synapses in layer 2/3 of the rat prefrontal cortex

**DOI:** 10.1101/2024.09.10.612266

**Authors:** Jiwoo Shin, Seung Yeon Lee, Yujin Kim, Suk-Ho Lee

**Affiliations:** Department of Brain and Cognitive Sciences, College of Natural Sciences, Seoul National University.; Department of Physiology, Seoul National University College of Medicine. 03080, Seoul, South Korea; Neuroscience Research Institute, Seoul National University College of Medicine, Seoul 03080, Republic of Korea

## Abstract

Short-term facilitation of recurrent excitatory synapses within the cortical network has been proposed to support persistent activity during working memory tasks, yet the underlying mechanisms remain poorly understood. We characterized short-term plasticity at the local excitatory synapses in layer 2/3 of the rat medial prefrontal cortex and studied its presynaptic mechanisms. Low-frequency stimulation induced slowly developing facilitation, whereas high-frequency stimulation initially induced strong depression followed by rapid facilitation. This non-monotonic delayed facilitation after a brief depression resulted from a high vesicular fusion probability and slow activation of Ca^2+^-dependent vesicle replenishment, which led to the overfilling of release sites beyond their basal occupancy. Pharmacological and gene knockdown (KD) experiments revealed that the facilitation was mediated by phospholipase C/diacylglycerol signaling and synaptotagmin 7 (Syt7). Notably, Syt7 KD abolished facilitation and slowed the refilling rate of vesicles with high fusion probability. Furthermore, Syt7 deficiency in layer 2/3 pyramidal neurons impaired the acquisition of trace fear memory and reduced c-Fos activity. In conclusion, Ca^2+^- and Syt7-dependent overfilling of release sites mediates synaptic facilitation at layer 2/3 recurrent excitatory synapses and contributes to temporal associative learning.

## Introduction

Synapses in the mammalian central nervous system undergo activity-dependent modulation of synaptic weight during repetitive stimulation, known as short-term synaptic plasticity (STP). STP is mediated by presynaptic mechanisms that regulate the release probability (p_r_), size of the readily releasable pool (RRP), and kinetics of vesicle replenishment of RRP (Zucker and Regehr, 2002; Jackman and Regehr, 2017; Neher and Brose, 2018). Short-term depression has been observed at synapses with a high baseline p_r_, primarily because of the rapid depletion of releasable vesicles during high-frequency stimulation (HFS) (Lin et al., 2022). Conversely, vesicle depletion is less pronounced at synapses with a low baseline p_r_, and cumulative increases in p_r_ during HFS are considered to mediate short-term facilitation (Rozov et al., 2001; Pan and Zucker, 2009; Aldahabi et al., 2024). Central synapses exhibit a wide spectrum of STP depending on different combinations of presynaptic factors mediating short-term depression and facilitation.

Previously, facilitation was modeled as an increase in p_r_ after an action potential (AP) and its slow decay during the inter-spike interval (ISI), resulting in a cumulative increase in p_r_ during HFS (Markram et al., 1998; Dittman et al., 2000). Recent studies have suggested that the number of docking (or release) sites in an active zone is limited and only partially occupied by releasable vesicles in the resting states (Miki et al., 2016; Pulido and Marty, 2018; Malagon et al., 2020; Lin et al., 2022). Therefore, p_r_ is determined by the vesicular fusion probability (p_v_) and release site occupancy (p_occ_). However, distinguishing the individual contributions of p_v_ and p_occ_ to p_r_ is challenging (Silva et al., 2021; Neher, 2024). It is also unclear whether facilitation is mediated by activity-dependent increases in the p_v_, p_occ_, or both.

Two recent models for vesicle priming and release view the ‘priming’ as a two-step reversible process that leads to a release-ready state. The release-ready vesicles could be vesicles in a tightly docked state (TS) or those occupying specialized docking sites (DS), and are supplied from vesicles in a loosely docked state (LS) or those occupying a distinct replacement sites (RS), which are called LS/TS model (Aldahabi et al., 2024; Neher, 2024) and RS/DS model (Miki et al., 2016; Silva et al., 2021), respectively. Common to both models is the assumption that release-ready vesicles constitute only a fraction of all docked vesicles under resting conditions and that this fraction is up- and down-regulated by synaptic activity.

Synaptotagmin-7 (Syt7) was recently found to play a crucial role in facilitation at different types of central synapses (Jackman et al., 2016; Chen et al., 2017; Martinetti et al., 2022). In addition to its role in facilitation, Syt7 accelerates refilling of the RRP after depletion and mediates asynchronous release (Bacaj et al., 2013; Liu et al., 2014; Jackman et al., 2016; Tawfik et al., 2021). Syt7-dependent synaptic facilitation is interpreted as an activity-dependent increase in p_v_ (Turecek et al., 2016; Jackman and Regehr, 2017; Turecek et al., 2017; Norman et al., 2023). However, it is unclear whether Syt7 contributes to facilitation through a Ca^2+^-dependent increase in p_occ_ (i.e., overfilling). Overfilling has been proposed as a mechanism supporting facilitation at the Drosophila neuromuscular junction (Kobbersmed et al., 2020), yet whether this process also occurs at the mammalian central synapses mediated by Syt7 has not been studied.

Each of layer 2/3 (L2/3) and layer 5 (L5) of neocortex displays intralaminar excitatory synapses between pyramidal cells comprising a recurrent network (Holmgren et al., 2003; Thomson and Lamy, 2007). Short-term facilitation at recurrent excitatory synapses in a cortical network may play a crucial role in maintaining working memory (Mongillo et al., 2008; Mongillo et al., 2012; Hansel and Mato, 2013). Although L2/3 pyramidal cells (PCs) represent task-relevant information during a delay period in working memory tasks (Fujisawa et al., 2008; Ozdemir et al., 2020), most studies on local excitatory recurrent synapses in the neocortex have focused on L5 exhibiting short-term depression (Reyes and Sakmann, 1999; Hempel et al., 2000; Yoon et al., 2020). Further, the synaptic properties of excitatory synapses in L2/3 remain poorly understood.

In the present study, we found that HFS of local excitatory synapses in L2/3 of the medial prefrontal cortex (mPFC) induced initial strong depression followed by delayed facilitation, whereas low-frequency stimulation induced slow monotonous facilitation. This unique form of STP could be explained by 1) high p_v_ of RRP vesicles at rest and throughout a train stimulation; 2) the low resting vesicular occupancy of RRP and its activity-dependent increase. These conditions led to delayed facilitation through progressive overfilling of release sites during HFS with high p_v_ vesicles beyond the resting occupancy. Syt7 knockdown (KD) in L2/3 PCs completely abolished these synaptic enhancements by inhibiting the Ca^2+^-dependent increase in release site occupancy. Finally, behavioral tests for trace fear conditioning showed that Syt7 KD impaired the acquisition of trace fear memory and reduced c-Fos expression in the mPFC. Collectively, these results support that Ca^2+^- and Syt7-dependent overfilling of release sites mediates synaptic facilitation at L2/3 recurrent excitatory synapses and contributes to temporal associative learning.

## Results

### Frequency dependence of STP at local excitatory synapses in L2/3 of the prelimbic cortex

We examined STP profiles at recurrent excitatory synapses between L2/3 PCs and at synapses from L2/3-PCs to fast-spiking interneurons (FSINs) in the prelimbic area of the rat mPFC. The PCs and FSINs were identified based on their distinct intrinsic properties and morphologies (***Figure 1—figure supplement 1***). To stimulate presynaptic PCs, L2/3 PCs were transfected with a plasmid encoding oChIEF using *in utero* electroporation (IUE; ***Figure 1A***; Lee et al., 2024). Photo-stimulation of the oChIEF-expressing cell soma at 40 Hz consistently evoked single APs with high fidelity throughout the 600-pulse trains (***Figure 1—figure supplement 2A-B***), which was consistent with little use-dependent inactivation of oChIEF (Lin et al., 2009; Hass and Glickfeld, 2016).

**Figure 1.**
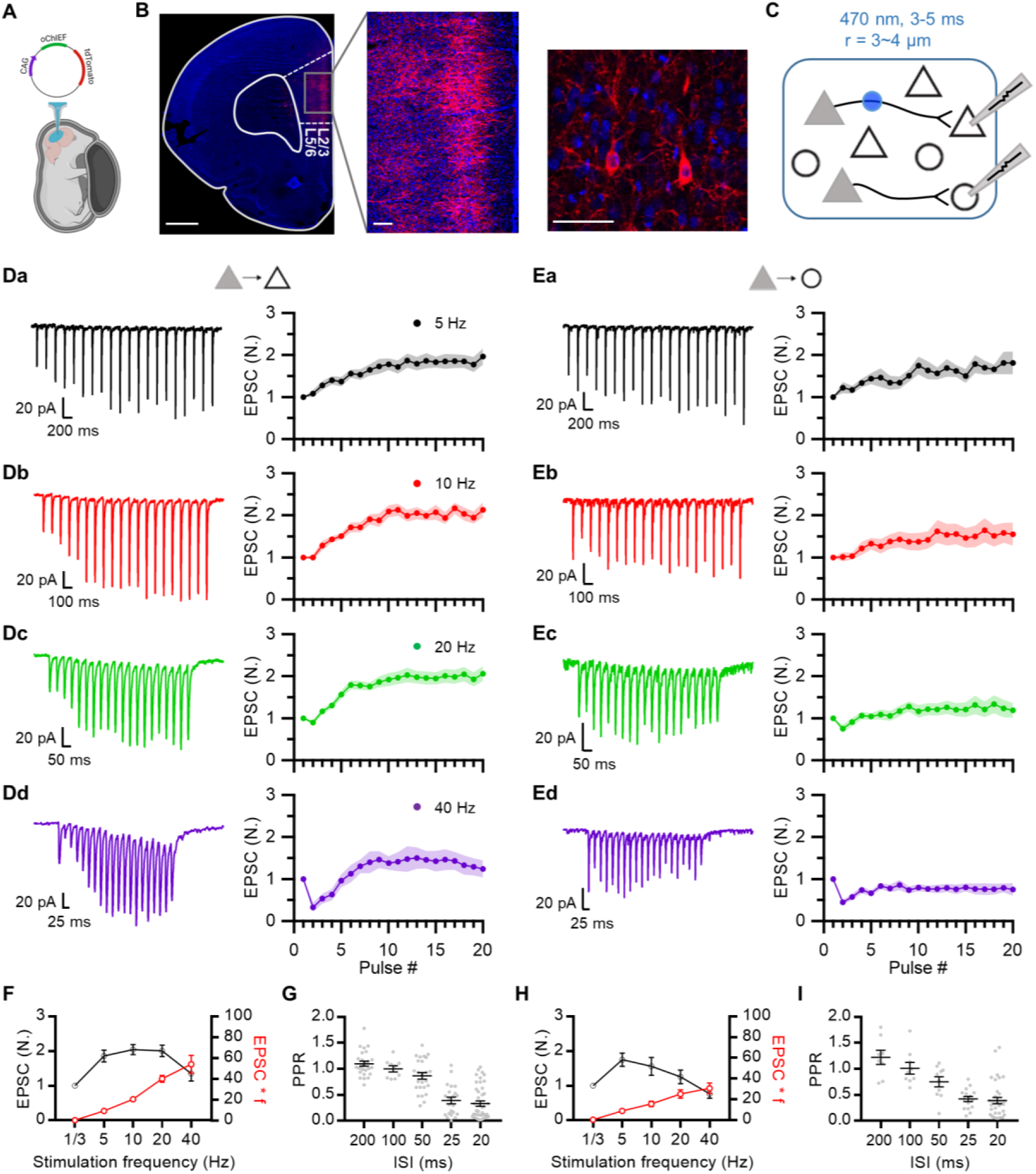
Frequency-dependence of short-term synaptic plasticity at prelimbic L2/3 excitatory synapses. (**A**) In utero electroporation following injection of the plasmid (CAG-oChIEF-tdTomato) into the ventricle of an embryo (E17.5). (**B**) Representative images showing specific expression of oChIEF-tdTomato in L2/3 pyramidal cells after IUE. The red fluorescence of tdTomato clearly visualizes oChIEF-expressing cell bodies in L2/3 and axons in L5. Scale bar: 1 mm, 100 μm, 50 μm from left to right. (**C**) Recording schematic showing photostimulation of oChIEF-expressing axon fibers of transfected PCs (*filled triangles*) and a whole-cell recording from non-transfected PCs (*empty triangles*) or FSINs (*empty circles*). A collimated DMD-coupled LED was used to confine the area of excitation (typically 3–4 μm in diameter) to a small region (*blue circle*) near the soma. (**D, E**) Representative traces for EPSCs averaged over 10 trials at each frequency (*left*) and average amplitudes of baseline-normalized EPSCs (*right*) during 20-pulse trains at frequencies from 5 to 40 Hz at PC-PC (*D*; n = 12, 9, 21, 10 cells for 5, 10, 20, 40 Hz, respectively) and PC-FSIN (*E*; n = 8, 9, 10, 10) synapses. Each data point was normalized to the average of the first EPSC. (**F**) Baseline-normalized amplitudes of steady-state EPSCs (EPSC_ss_; *black symbols*) and synaptic efficacy (EPSC_ss_ × *f*; *red symbols*) as a function of stimulation frequency (*f*) at PC–PC synapses. EPSC_ss_ was measured from the average of last 5 EPSCs from the 20-pulse trains (n = 12, 9, 21, 10). (**G**) Paired pulse ratio (PPR) as a function of inter-spike intervals (n = 25, 9, 25, 22, 43 for 200, 100, 50, 25, 20 ms ISI, respectively) at PC-PC synapses. (**H**) EPSC_ss_ and synaptic efficacy at PC-FSIN (n = 8, 9, 10, 10). (**I**) PPR at PC-FSIN (n = 8, 9, 10, 15, 34). *Gray symbols*, individual data.

Given that IUE transfected approximately 10-20% of L2/3 PCs (***Figure 1A***), we made a whole-cell patch on a non-transfected L2/3 PC or an FSIN (***Figure 1C***). To stimulate a minimal number of excitatory axon fibers, EPSCs were evoked using minimal optical stimulation (***Figure 1—figure supplement 3A***; 470 nm, 3-4 μm in radius and 3-5 ms in duration). EPSCs evoked by minimal stimulation were examined to determine the uniformity of the kinetic properties (***Figure 1—figure supplement 3B-C***). Moreover, we reassessed the kinetics of EPSCs during trains *post hoc* to ensure uniformity. EPSCs were evoked by 20-pulse trains at 4 different frequencies (5, 10, 20, and 40 Hz) to cover the range of firing rates of PCs observed *in vivo* during working memory tasks (Baeg et al., 2003). Excitatory synapses from PCs onto both other PCs and FSINs (PC–PC and PC–FSIN) showed strong (∼2-fold enhancement) and lasting (several seconds) short-term facilitation at 20 Hz and lower frequencies (***Figure 1D-E***). The PC-PC synapses exhibited stronger facilitation than did the PC-FSIN synapses.

Steady-state EPSCs at the PC-PC synapses were not significantly different from 5 to 20 Hz (***Figure 1F***). Frequency invariance at the PC-PC synapses suggests that Ca^2+^-dependent synaptic facilitation may counteract vesicle depletion in a frequency-dependent manner (Turecek et al., 2017; Lin et al., 2022). Despite facilitation at all frequencies, the paired pulse ratio (PPR) rapidly decreased as the ISI decreased (***Figure 1G, I***). At 40 Hz stimulation, both PC-PC and PC-FSIN synapses initially underwent strong depression then followed by facilitation in a few stimuli (***Figure 1Dd, Ed***). Such delayed facilitation after paired pulse depression (PPD) has not been previously observed in central synapses except for a slight increase in EPSC at the cerebellar synapses under artificially depolarized conditions (Pulido and Marty, 2018).

We confirmed that optically evoked EPSCs were strictly dependent on AP generation (***Figure 1—figure supplement 4A***), suggesting that EPSCs are evoked by optical stimulation of presynaptic axon fibers rather than by direct depolarization of presynaptic boutons. Additionally, dual patch-clamp techniques showed that the STP of optically evoked EPSCs did not differ from the electrically evoked STP of EPSCs at the PC-FSIN synapses (***Figure 1—figure supplement 4B***). Finally, we confirmed that AMPA receptor (AMPAR) desensitization was not responsible for PPD at 40 Hz, ruling out the possible effects of postsynaptic factors on STP (***Figure 1—figure supplement 5A***), and that AMPAR saturation was not significant during facilitation (***Figure 1—figure supplement 5B***).

### Delayed facilitation results from slow activation of Ca^2+^-dependent vesicle replenishment at a constantly high vesicular fusion probability

The strong PPD at 40 Hz suggests a high vesicular fusion probability (p_v_) and likely tight coupling of the releasable vesicles to a Ca^2+^ source. To test this hypothesis, we examined the effects of EGTA, a slow calcium chelator, on synaptic transmission at the PC-PC synapses. The baseline EPSC amplitude (EPSC_1_) was not affected by incubation of the slice in the bath solution containing 50 μM EGTA-AM for longer than 20 min, supporting tight coupling. Moreover, the PPR was decreased to 0.19, and subsequent facilitation slowed but was still distinct (***Figure 2A-B***). These findings indicate that the initial p_v_ should be higher than 0.81 considering vesicle recruitment during ISI. The effects of EGTA-AM were confirmed in the same cell by measuring the baseline EPSC and PPR before and after applying EGTA-AM to the bath. EPSC_1_ remained unchanged, while PPR was reduced by half (***Figure 2—figure supplement 1***).

**Figure 2.**
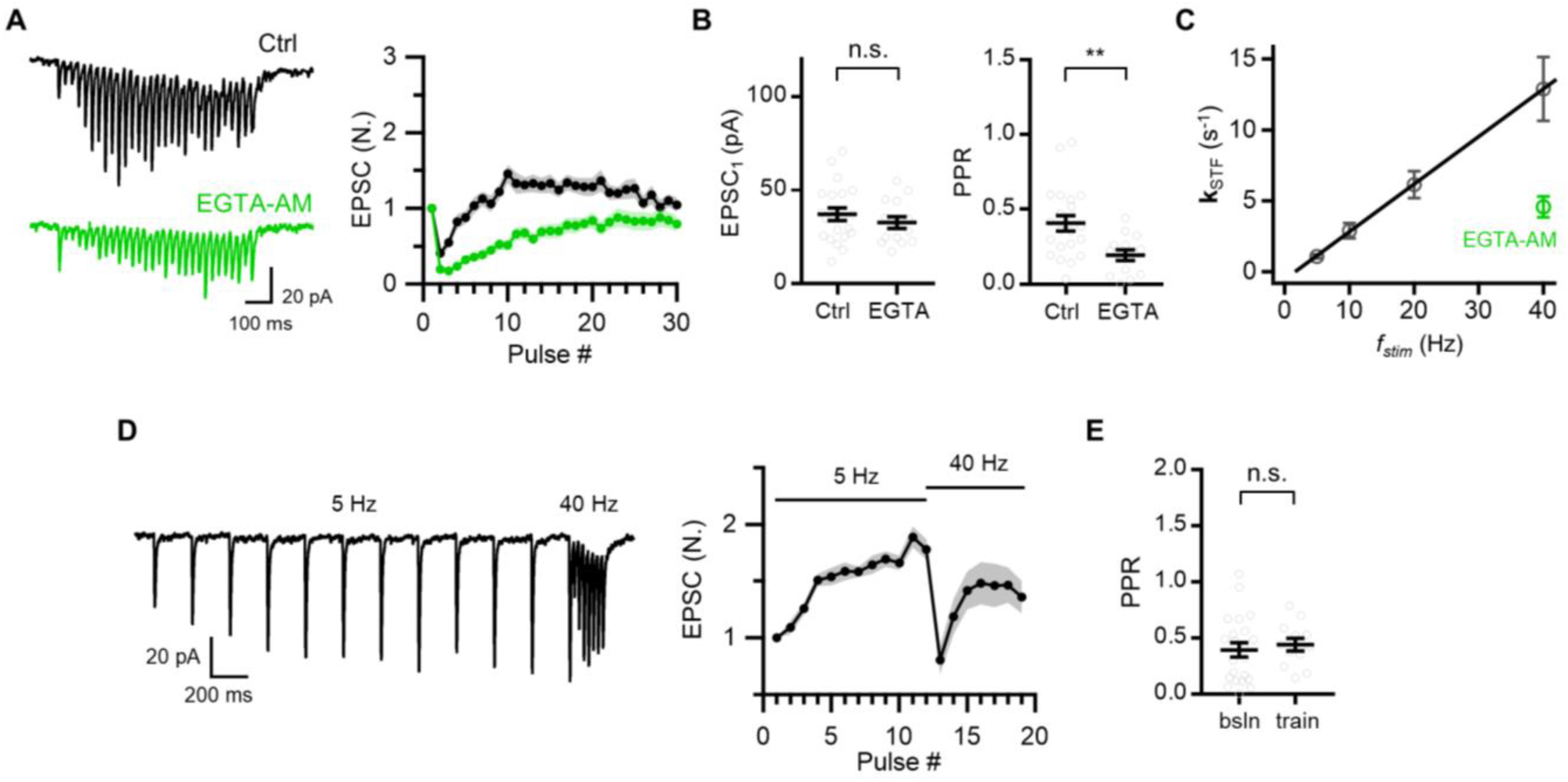
Delayed facilitation results from slow activation of Ca^2+^-dependent vesicle replenishment at a constantly high vesicular fusion probability. (**A**) Representative traces (*left*) and mean baseline-normalized amplitudes (*right*) of EPSCs evoked by 30-pulse trains at 40 Hz in control (n = 21; *black*) and in the presence of 50 μM EGTA-AM (n = 14; *green*). (**B**) Mean values for the first EPSC amplitude (EPSC_1_, *left*) and PPR (*right*) from the experiments displayed in (A). *Gray symbols*, individual data. (**C**) Plot of rate constants for short-term facilitation (**k**_STF_) as a function of stimulation frequency (*f_stim_*), showing a linear relationship. The linear regression line (*black*) is shown fitted to **k**_STF_ values, estimated from Figure 1. (**D**) Representative EPSC traces (*left*) and average of baseline-normalized EPSCs (*right*) evoked by 12-pulse stimulation at 5 Hz, followed by 40 Hz 7-pulse train (n = 12). Note that slowly developing facilitation was converted to rapid facilitation after strong PPD. (**E**) Mean values for PPR at 40 Hz. The baseline PPR was reproduced from Figure 1G and the PPR during 5 Hz train was calculated as (13^th^ EPSC) / (12^th^ EPSC). *Gray symbols*, individual data. All statistical data are represented as mean ± S.E.M.; n.s. = not significant; **, P<0.01; unpaired t-test.

Notably, the application of EGTA-AM markedly slowed facilitation at 40 Hz (***Figure 2A, C***). The frequency invariance of steady-state EPSCs (***Figure 1F***) and the slower facilitation at 40 Hz in the presence of EGTA (***Figure 2A***) imply a Ca^2+^-dependent increase in the refilling rate of the RRP and/or an increase in the p_v_ of reluctant vesicles in light of previous studies (Hosoi et al., 2007; Liu et al., 2014; Ritzau-Jost et al., 2014; Turecek et al., 2016; Ritzau-Jost et al., 2018; Kusick et al., 2020). Moreover, facilitation was accelerated at higher stimulation frequencies. The plot of the rate constant for the increase in EPSCs (denoted as **k**_STF_, the rate constant for the development of short-term facilitation) as a function of the stimulation frequency (*f_stim_*) revealed a linear relationship (***Figure 2C***). **k**_STF_ in the presence of EGTA-AM was distinctly below this relationship, supporting the Ca^2+^-dependent acceleration of facilitation.

**k**_STF_ was also accelerated when the stimulation frequency was increased from 5 Hz to 40 Hz (***Figure 2D***; 1.27 ± 0.18/s to 24.27 ± 3.44/s). Notably, the second 40-Hz train stimulation led to strong PPD, followed by accelerated facilitation (***Figure 2D-E***). The strong PPD suggests that the vesicular fusion probability kept high during the 5-Hz stimulation, arguing against the possibility for an increase in p_v_. Both two-step sequential priming models, LS/TS and RS/DS models, are suitable to interpret our data and we will refer to the different populations of vesicles as TS and LS vesicles, respectively (Taschenberger et al., 2016; Neher and Brose, 2018; Lin et al., 2022; Neher, 2024). Within this framework, vesicles are released from TS with high p_v_ both in the resting state and during facilitation, and the delayed facilitation is seen as a result of Ca^2+^-dependent progressive increase in the occupancy of TS vesicles on docking sites.

### Presynaptic calcium measurements at axonal boutons of L2/3 pyramidal cells

To test contribution of Ca^2+^ channel regulation to STP (Dobrunz et al., 1997; Jackman and Regehr, 2017), we estimated total amount of Ca^2+^ increments during 40 Hz AP trains at axonal boutons of L2/3 PCs in the mPFC. To this end, we measured AP-evoked Ca^2+^ transients (AP-CaTs) using two-dye ratiometry techniques, and estimated calcium binding ratio from AP-CaTs (***Figure 2—figure supplement 2A-C***). Measuring the amplitudes and decay time constants of AP-CaTs at different concentrations of Fluo-5F (150, 250 or 500 μM), and plotting them as a function of calcium binding ratio of Fluo-5F (κ_B_), we estimated the amplitude of AP-CaTs (A_Ca_ = 1.16 μM), Ca^2+^ decay time constant (τ_c_ = 43 ms) in the absence exogenous Ca^2+^ buffer, and endogenous static Ca^2+^ binding ratio (κ_S_ = 96.4 or 77.3 estimated from ***Figure 2—figure supplement 2D-E***, respectively; *See Materials and Methods*). The resting [Ca^2+^] ([Ca^2+^]_rest_) was measured as 50 nM (***Figure 2—figure supplement 2F***). Estimating the total Ca^2+^ increments for the first four AP-CaTs in a 40 Hz train according to Equation 2 (*Materials and Methods*), we found no evidence for inactivation or facilitation of calcium influx during a 40 Hz train (***Figure 2—figure supplement 2G-I***), arguing against the possibility that Ca^2+^ channel inactivation and facilitation contribute to strong PPD and delayed facilitation during a 40 Hz train.

### Release sites have low baseline occupancy and these are increased during facilitation and post-tetanic augmentation

Given that p_v_ was greater than 0.8, the two-fold increase in EPSC during 20 pulse train stimulations could not be explained by an increase in p_v_. Recent studies have shown that vesicle release sites (N) are limited in number and are partially occupied at rest (Miki et al., 2016; Pulido and Marty, 2018; Malagon et al., 2020; Lin et al., 2022). Under such a high p_v_, two-fold synaptic facilitation should be attributed to an increase in p_occ_. The vesicle replenishment rate is accelerated during HFS in a Ca^2+^-dependent manner (***Figure 2A-C***; Hosoi et al., 2007). This may lead to the overfilling of release sites beyond their basal vesicular occupancy, which in turn may increase EPSC under high p_v_. To test this hypothesis, we determined the quantal parameters at the L2/3 local excitatory synapses. We first measured the quantal size from asynchronous release events after replacing extracellular Ca^2+^ with Sr^2+^, as previously described (Yoon et al., 2020) (***Figure 3—figure supplement 1A-B***). The mean and coefficient of variance of the quantal sizes were 24 pA and 0.25 at PC-PC synapses, respectively, and were 30 pA and 0.31 at PC-FSIN synapses (***Figure 3—figure supplement 1C***).

Next, we performed a variance-mean (V-M) analysis of phasic EPSCs at excitatory synapses during train stimulation at 5 Hz and 40 Hz (Scheuss et al., 2002). Applying multiple-probability fluctuation analysis (MPFA) using intra-site quantal variability of 0.2 (Valera et al., 2012), the V-M plot was best fitted by the number of release sites (N) of 5.3 and the quantal size (q) of 25 pA at the PC-PC synapses (***Figure 3A***). The p_r_ for EPSC_1_ was calculated as 0.32. We performed the same analysis for the PC-FSIN synapses and found N = 5.3, q = 35 pA, and p_r_ for EPSC_1_ = 0.31 (***Figure 3B***). Our estimate for N was smaller but comparable to the estimate (6.9) at excitatory synapses in L2/3 of the mouse barrel cortex (Holler et al., 2021).

**Figure 3.**
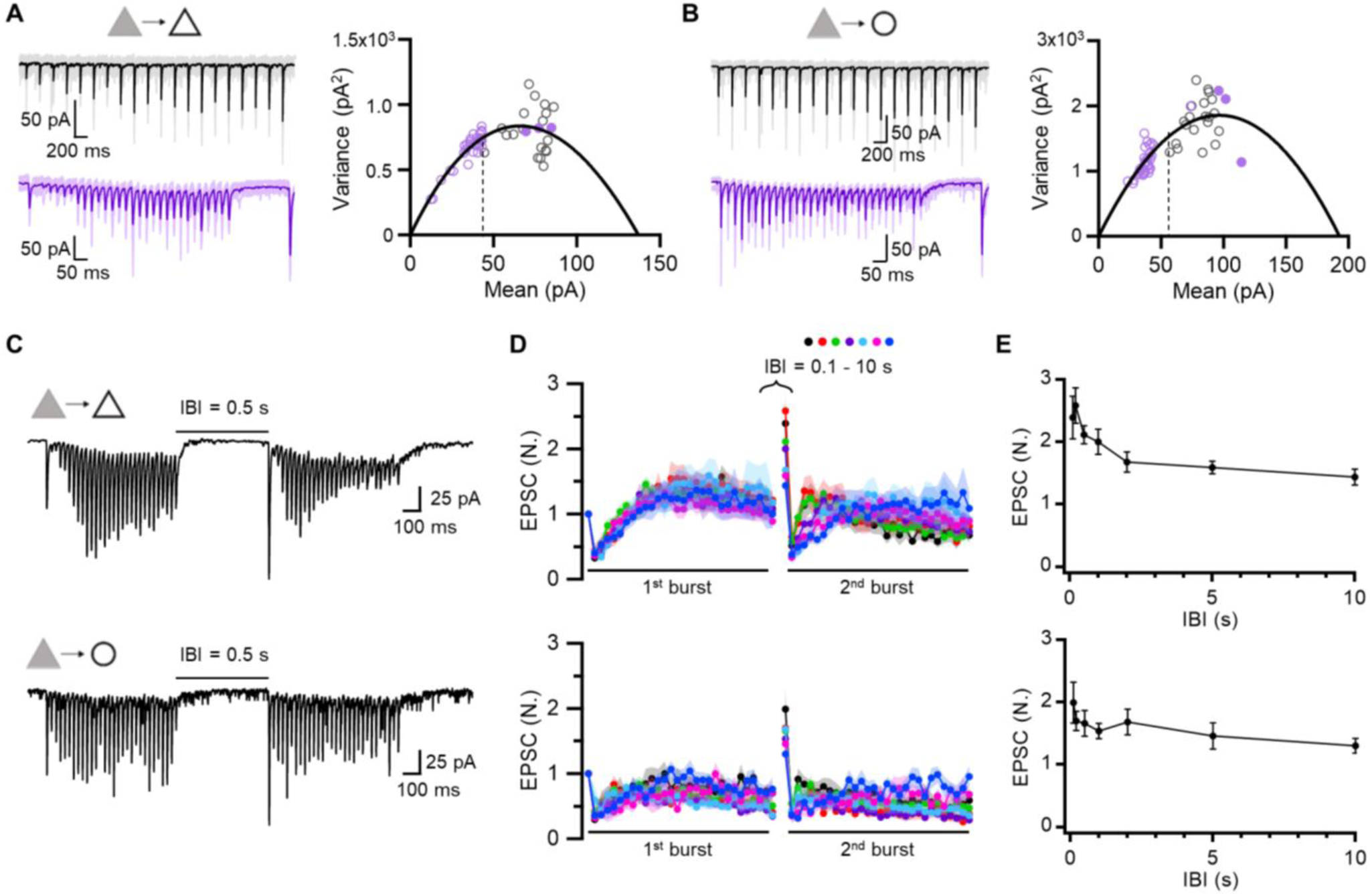
Low baseline occupancy of release sites and its increase during facilitation and post-tetanic augmentation. (**A, B**) *Left*, Representative EPSCs evoked by 5 and 40 Hz train stimulation (*black*, 5 Hz; *purple*, 40 Hz). *Right*, Variance-mean plots of EPSCs amplitude from averaged EPSCs recorded at PC-PC (A; n = 12, 17 for 5 and 40 Hz, respectively) and PC-FSIN (B; n = 8, 10) synapses. The data were fitted using multiple-probability fluctuation analysis (MPFA). Error bars are omitted for clarity. The 1^st^ EPSC of 5 Hz train (*broken line*) was used to estimate the resting level of p_occ_. *Filled circles* were measured from post-tetanic augmented EPSCs (*A*, n = 12, 12, 9; *B*, n = 7, 7, 7 for 0.1, 0.2, 0.5 s IBIs, respectively). (**C-E**) Post-tetanic augmentation (PTA) experiments at PC-PC (*top*) and PC-FSIN (*bottom*) synapses. (**C**) Representative traces for EPSCs evoked by double 40 Hz train stimulations separated by 0.5 s. (**D**) Mean baseline-normalized amplitudes of EPSCs evoked by double 40 Hz trains at different inter-burst intervals (IBIs, 0.1, 0.2, 0.5, 1, 2, 5, 10 s). *Upper*, PC-PC synapse (n = 12, 12, 21, 11, 11, 16, 11 from short to long IBIs, respectively). *Lower*, PC-FSIN synapse (n = 10, 9, 8, 9, 9, 7, 9). (**E**) PTA time course. The baseline-normalized amplitudes of 1^st^ EPSC from the 2^nd^ train were plotted as a function of IBIs.

The p_r_ estimate was clearly smaller than the possible minimum value for p_v_ (0.81) estimated from the PPR data (***Figure 2B***). Given that p_r_ is a product of p_occ_ and p_v_ (Neher, 2024), a p_r_ smaller than p_v_ indicates the partial occupancy of release sites in the resting state. If synaptic facilitation were mediated by the recruitment of new release sites, it would be accompanied by an increase in the variance of augmented EPSCs (Valera et al., 2012; Kobbersmed et al., 2020). However, the augmented EPSCs during facilitation followed the same parabola in the V-M plot (***Figure 3A-B***). Therefore, facilitation appears to be mediated by the progressive overfilling of a finite number of release sites beyond the basal occupancy. In the next section, we show that p_v_ is close to unity based on high double failure rate upon paired pulse stimulation (***Figure 3—figure supplement 2***) and little effect of extracellular [Ca^2+^] ([Ca^2+^]_o_) elevation on baseline EPSC amplitudes (***Figure 3—figure supplement 3***). Assuming this, the baseline p_r_ at the first pulse estimated from the fitted parabolic curve can be largely regarded as a baseline p_occ_ (c.a. 30%), and the two-fold increase in EPSCs during facilitation can be attributed to an increase of up to 60% in p_occ_.

We investigated the time course of recovery from facilitation and post-tetanic augmentation (PTA) by applying dual-train stimulations (30 pulses at 40 Hz) separated by different inter-burst intervals (IBIs; ***Figure 3C-D***). The initial EPSCs in the second train were enhanced by more than twofold, and followed by a strong PPD, suggesting that fusion probability stays high in the IBIs and thus synaptic transmission is carried by TS vesicles. The decay time course of PTA was characterized by two phases (initially fast and later slow; ***Figure 3E***), which may reflect a two-phase Ca^2+^ decay or two distinct recovery processes that occur sequentially. The variance of potentiated EPSCs measured at IBIs of 0.1, 0.2, and 0.5 s was not different from that of EPSCs at the peak of facilitation (filled circles, ***Figure 3A-B***). Given that the same RRP mediates both facilitation and augmentation and that post-tetanic EPSCs is carried by TS vesicles similar to the baseline EPSCs, this plot suggests that PTA, similar to short-term facilitation, is mediated by an increase in the TS vesicle occupancy.

### High double failure rate and little effect of [Ca^2+^]_o_ elevation on baseline EPSC amplitudes suggest that vesicular fusion probability is close to one

The vesicle dynamics of STP has been explained using a simple refilling model (Hosoi et al., 2007; Neher and Sakaba, 2008). This model assumes that the vesicles are reversibly docked to a limited number of release sites (N) with forward and reverse rate constants (denoted as k_1_ and b_1_, respectively). Under the framework of the simple refilling model (see *Materials and Methods*), the rate constant for the refilling of the RRP after depletion by the first pulse is the sum of the baseline k_1_ and b_1_. This is estimated as 23/s from the plot of PPR as a function of ISIs (***Figure 3—figure supplement 2A***). From the baseline occupancy of 0.3 and p_occ_ = k_1_ / (k_1_ + b_1_), we estimated k_1_ to be 6.9/s.

When paired APs were applied with an ISI of 20 ms, the failure rate at 1^st^ pulse, P(F_1_), was 10.6%, and the probability of two consecutive failures, P(F_1_, F_2_), was 6.2% (***Figure 3—figure supplement 2A***). From the equation P(F_1_) = (1 – p_r_)^N^, p_r_ was calculated as 0.312 when N = 6, comparable to p_r_ (= 0.32) estimated from the V-M analysis (***Figure 3A***). We calculated P(F_1_, F_2_) under the framework of a simple refilling model, with p_v_ and k_1_ set as free parameters and other parameters set according to the following relationships: P(F_1_) = (1 – p_r_)^N^, p_r_ = p_v_·p_occ_, and p_occ_ = k_1_ / (k_1_ + b_1_). The calculated values for P(F_1_, F_2_) and their difference from the observed value (6.2%) are shown in the plane of k_1_ *vs*. p_v_ in ***Figure 3—figure supplement 2B-C***. Given that k_1_ was greater than 5/s, the difference between the calculated and observed values of P(F_1_,F_2_) was minimal when p_v_ = 1 (***Figure 3—figure supplement 2C***).

To find optimal values for k_1_ and p_v_ that best explain observed probabilities shown in ***Figure 3—figure supplement 2Ac***, a cost function was implemented as the sum of squared errors between observed and predicted probability values for the four combinations of success and failure of 1st and 2nd EPSCs as described in *Materials and Methods*. The minimum of the cost function (5.63 x 10^-5^) was found at k_1_ = 5.21/s and p_v_ = 0.999, which predicted P_11_ = 41.5%, P_10_ = 47.9%, P_01_ = 4.93%, and P_00_ = 5.67% (subscript 1 and 0 denote success and failure, respectively).

To further validate that baseline p_v_ is near to unity, we examined EPSC changes following elevation of [Ca²⁺]ₒ from 1.3 to 2.5 mM. According to the fourth-power relationship between synaptic responses and [Ca²⁺]ₒ (Equation 3 in (Dodge Jr and Rahamimoff, 1967)), a 3.24-fold increase in the EPSC amplitude was expected. However, the EPSC amplitude increased only 1.23-fold in average, a change that was not statistically significant (***Figure 3—figure supplement 3A-B***). This small response suggests that p_v_ is already saturated at rest, limiting the dynamic range for further enhancement through increased calcium influx. Recent morphological and functional studies revealed that elevation of [Ca^2+^]_o_ induces an increase in the number of TS or docked vesicles to a similar extent as our observation (Kusick et al., 2020; Lin et al., 2025), raising a possibility that an increase in p_occ_ is responsible for the 1.23-fold increase in EPSC at high [Ca^2+^]_o_. A slight but significant reduction in PPR was observed under high [Ca²⁺]ₒ too. An increase in p_occ_ is thought to be associated with that in the baseline vesicle refilling rate. While PPR is always reduced by an increase in p_v_, the effects of refilling rate to PPR is complicated. For example, PPR can be reduced by both a decrease (***Figure 2—figure supplement 1***) and an increase (Lin et al., 2025) in the refilling rate induced by EGTA-AM and PDBu, respectively. Thus, the slight reduction in PPR is not contradictory to the possible contribution of p_occ_ to the high [Ca^2+^]_o_ effects. Notably, when slices were pre-incubated with 50 μM EGTA-AM, elevating extracellular [Ca^2+^] from 1.3 to 2.5 mM produced no significant change in either baseline EPSC amplitude or PPR (***Figure 3—figure supplement 3C-D***), supporting that the modest Ca^2+^-dependence of baseline EPSCs and PPR in the absence of EGTA is primarily mediated by a change in p_occ_ rather than p_v_.

### Pharmacological manipulations reveal specific molecular identities underlying vesicle loading processes

Our results suggest that the baseline occupancy of TS vesicles is low and that the activity-dependent progressive overfilling of docking sites with TS vesicles is responsible for short-term facilitation and augmentation. To elucidate the specific molecular link between synaptic enhancement and Ca^2+^-dependent overfilling, we examined the effects of several drugs known to affect vesicle dynamics. We first examined the roles of phospholipase C (PLC) and diacylglycerol (DAG) by applying 5 μM U73122, a PLC inhibitor, or 20 μM 1-oleoyl-2-acetyl-sn-glycerol (OAG), a DAG analog. The STP at 40 Hz and PTA at 0.5-s interval were examined before and after applying each drug to the bath. For both drugs, the facilitation and augmentation were reduced (***Figure 4A-B***). U73122 did not affect the basal EPSC amplitude (EPSC_1_) but resulted in a stronger PPD, slower facilitation, and lower augmentation (***Figure 4A***). These effects of U73122 suggest the involvement of Ca^2+^-induced PLC activation in progressive overfilling. In contrast, OAG increased EPSC_1_ while maintaining a pronounced PPD (***Figure 4B***), indicating an increase in the TS vesicle pool size. Meanwhile, OAG slowed **k**_STF_ and reduced augmentation, indicating occlusion of activity-dependent synaptic facilitation. Phorbol esters and DAG are thought to accelerate priming through Munc13 activation (Rhee et al., 2002; Taschenberger et al., 2016; Aldahabi et al., 2022); they also shorten the bridges that interlink docked vesicles with the active zone (AZ) membrane (Papantoniou et al., 2023). Collectively, these suggest that PLC- and DAG-mediated signaling play a role in the overfilling of docking sites underlying short-term facilitation and augmentation.

**Figure 4.**
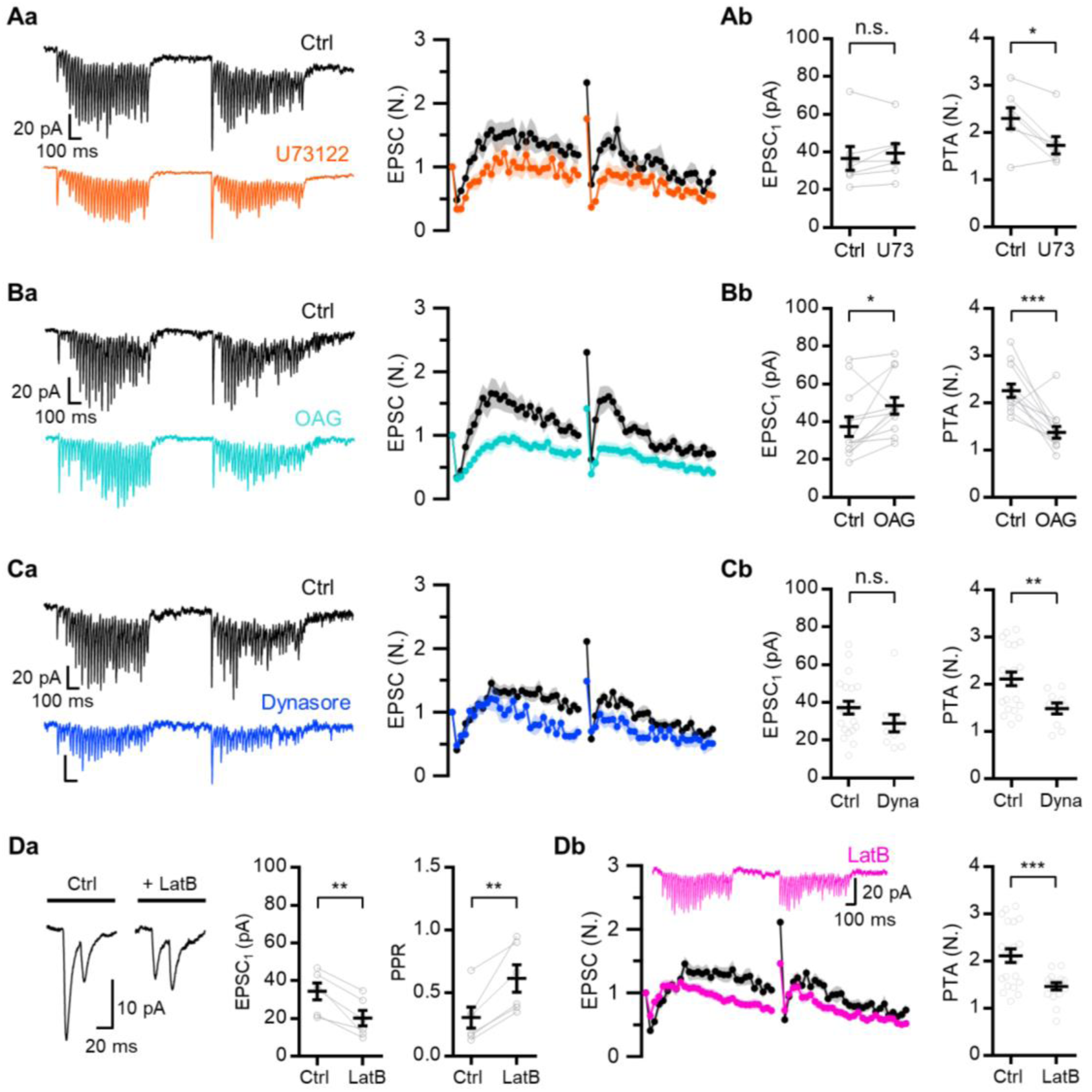
Pharmacological experiments reveal specific molecular mechanisms underlying vesicle loading processes. (**A-C**) (**a**) Representative EPSC traces (*left*) and mean baseline-normalized EPSCs (*right*) evoked by double 40 Hz train stimulations separated by 0.5 s inter-burst interval (IBI) in control and in the presence of 5 μM U73122 (*Aa*, n = 7, *orange*), 20 μM OAG (*Ba*, n = 12, *cyan*) or 100 μM dynasore (*Ca*, n = 10, *blue*). (**b**) Mean values for baseline EPSCs (EPSC_1_, *left*) and augmentation (*right*) from the experiments shown in corresponding *a* panel. (**Da**) Representative EPSC traces evoked by paired pulses (*left*) and mean values for baseline EPSC amplitude (*middle*) and PPR (*right*) before and after applying 20 μM LatB (n = 6). (**Db**) Representative EPSC traces (*left, upper*) and average of normalized EPSCs (*left, lower*) evoked by double 40 Hz train stimulation separated by 0.5 s in control (n = 21) and in 20 μM LatB conditions (n = 16; *pink*). *Right*, Mean values for augmentation in control and LatB conditions. *Gray symbols*, individual data. All statistical data are represented as mean ± S.E.M., *, P<0.05; **, P<0.01; ***, P<0.001; unpaired or paired t-test; n.s. = not significant.

Although the acceleration of Ca^2+^-dependent vesicle refilling supports rapid facilitation at 40 Hz, the later part of the delayed facilitation underwent a slight depression, which was more pronounced during the second burst (***Figure 3D* *and* *Figure 4***). Sustained synaptic transmission is limited by the spatial availability of docking sites because it is undermined by accumulation of exocytic remainings in the presynaptic AZ during HFS (Neher, 2010; Haucke et al., 2011). To verify this, we examined the effects of dynasore (100 μM), a specific inhibitor of endocytosis (Macia et al., 2006). The treatment with dynasore did not influence EPSC_1_ or the early phase of facilitation, but it exacerbated depression during the later phase and attenuated PTA (***Figure 4C***). This suggests that late synaptic transmission during the 40-Hz train is limited by site clearance.

Given the involvement of actin polymerization in the multiple steps of vesicle docking and recycling, we examined the effects of 20 μM latrunculin B (LatB), an inhibitor of actin polymerization. The effects of LatB were complicated: (1) EPSC_1_ was decreased, whereas PPR was increased (***Figure 4Da***) and (2) facilitation and augmentation were reduced (***Figure 4Db***). The latter effect may result from defects in vesicle replenishment rates, either through the impaired physical movement of vesicles or disrupted site clearance (Sakaba and Neher, 2003; Lee et al., 2012; Hallermann and Silver, 2013; Miki et al., 2016), as Ca^2+^-dependent vesicle refilling mediates facilitation and augmentation (***Figure 2* *and* *Figure 3***). Meanwhile, the former effect might be attributed to a shift in equilibrium in the priming states towards a loosely docked state (LS) at rest. It is unknown whether actin affects vesicular fusogenicity in neurons, although it facilitates multiple steps of vesicle fusion by enhancing membrane tension in neuroendocrine cells (Wu and Chan, 2022).

### Short-term facilitation at both types of local excitatory synapses in L2/3 is abolished by Syt7 knockdown

We examined whether Syt7 mediates facilitation at the PC-PC and PC-FSIN synapses in L2/3 of the mPFC. To this end, oChIEF and shRNA against Syt7 (shSyt7) were co-expressed in L2/3 PCs using IUE. Syt7 KD resulted in a complete loss of facilitation at all tested stimulation frequencies (***Figure 5A-B***). Short-term depression and paired-pulse depression were more pronounced as the stimulation frequency increased, and the frequency invariance of steady-state EPSC was abolished, implying little contribution of Ca^2+^-dependent vesicle recruitment (***Figure 5C-F***). The lack of facilitation could not be attributed to the failure of APs in presynaptic axons, as optical stimulation elicited reliable APs in Syt7 KD PCs during the 40-Hz train (***Figure 1—figure supplement 2C***). Syt7 KD had little effect on the basal properties, including the peak amplitude, rise/decay time, and time-to-peak, of single AP-induced EPSCs (***Figure 1—figure supplement 3D-E***). Moreover, Syt7 KD did not significantly affect the intrinsic properties of neurons (***Figure 5—figure supplement 1***).

**Figure 5.**
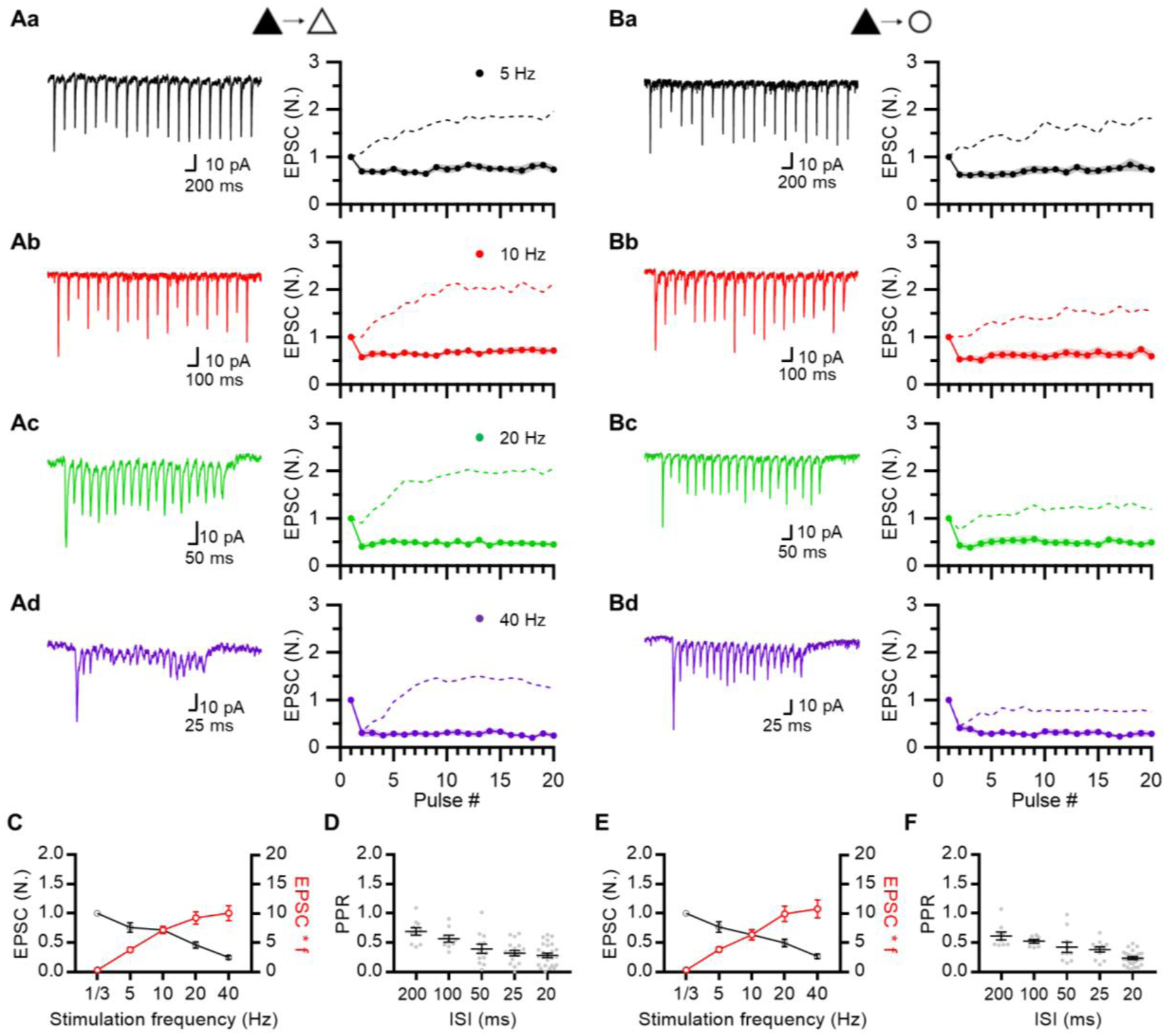
STF at both types of local excitatory synapses is abolished by Syt7 KD. (**A, B**) Representative EPSC traces (*left*) and mean baseline-normalized amplitudes of EPSCs (*right*) evoked by 20-pulse trains at 5 to 40 Hz. STP was measured at PC-PC (*A*; n = 10, 9, 12, 9) and PC-FSIN (*B*; n = 9, 8, 10, 9) synapses, in which presynaptic Syt7 transcripts were depleted (Syt7 KD). Syt7 KD pyramidal cells are indicated as *black triangles* on the top. For comparison, STP in WT synapses is reproduced from Figure 1 (*dotted lines*). Same frequency color codes were used as in Figure 1. (**C**) Baseline-normalized amplitudes of steady-state EPSC (EPSC_ss_; *black symbols*) and synaptic efficacy (EPSC_ss_ × *f*; *red symbols*) as a function of stimulation frequency (*f*) at PC-PC synapses. EPSC_ss_ was defined as the average of last 5 EPSC amplitudes from 20-pulse trains. (**D**) PPR as a function of inter-spike intervals (n = 10, 9, 12, 18, 26). (**E**) EPSC_ss_ and synaptic efficacy at PC-FSIN synapses. (**F**) PPR at PC-FSIN synapses (n = 9, 8, 10, 12, 24). *Gray symbols*, individual data.

To examine whether Syt7 KD had any nonautonomous effects on postsynaptic cells, we measured the quantal size (q) from asynchronous events in the presence of Sr^2+^ and found that the mean quantal size remained unaltered at Syt7 KD synapses (***Figure 3—figure supplement 1C***). This indicated that Syt7 did not influence postsynaptic parameters, including the density and kinetic properties of AMPARs. To test the specificity of shSyt7, we examined whether co-expression of the shRNA-resistant form of Syt7 restored synaptic facilitation in both excitatory synapse types. Presynaptic overexpression of shRNA-resistant Syt7 under the CAG promoter rescued the effect of shRNA at both PC-PC and PC-FSIN synapses (***Figure 5—figure supplement 2***). These results underscore the crucial role of presynaptic Syt7 in mediating short-term facilitation at local excitatory synapses in L2/3 of the mPFC.

### Syt7 KD synapses exhibit complementary changes in the number of release sites and their vesicle occupancy

To estimate the quantal parameters in Syt7 KD synapses, we performed V-M analysis of EPSCs evoked by train stimulation (***Figure 6A-B***). Fitting a parabola to the V-M plot revealed higher baseline p_occ_ in Syt7 KD synapses than in wild-type (WT) synapses for both synapse types (KD, 0.68 and 0.42; WT, 0.32 and 0.31 for PC-PC and PC-FSIN synapses, respectively). The p_occ_ monotonously decreased during a train stimulation, as expected from short-term depression. Moreover, the number of release sites (N) was markedly lower in Syt7 KD synapses (N = 3.5) than in WT synapses (N = 5.3; ***Figure 3A***). Next, we examined the PTA induced by the same protocol as in ***Figure 3*** (two 40-Hz train stimulations separated by variable IBIs) at Syt7 KD synapses (***Figure 6C***). The 40 Hz train-induced synaptic depression quickly recovered during IBIs, but no PTA was observed at Syt7 KD synapses, indicating its crucial role in augmentation (***Figure 6D-E***).

**Figure 6.**
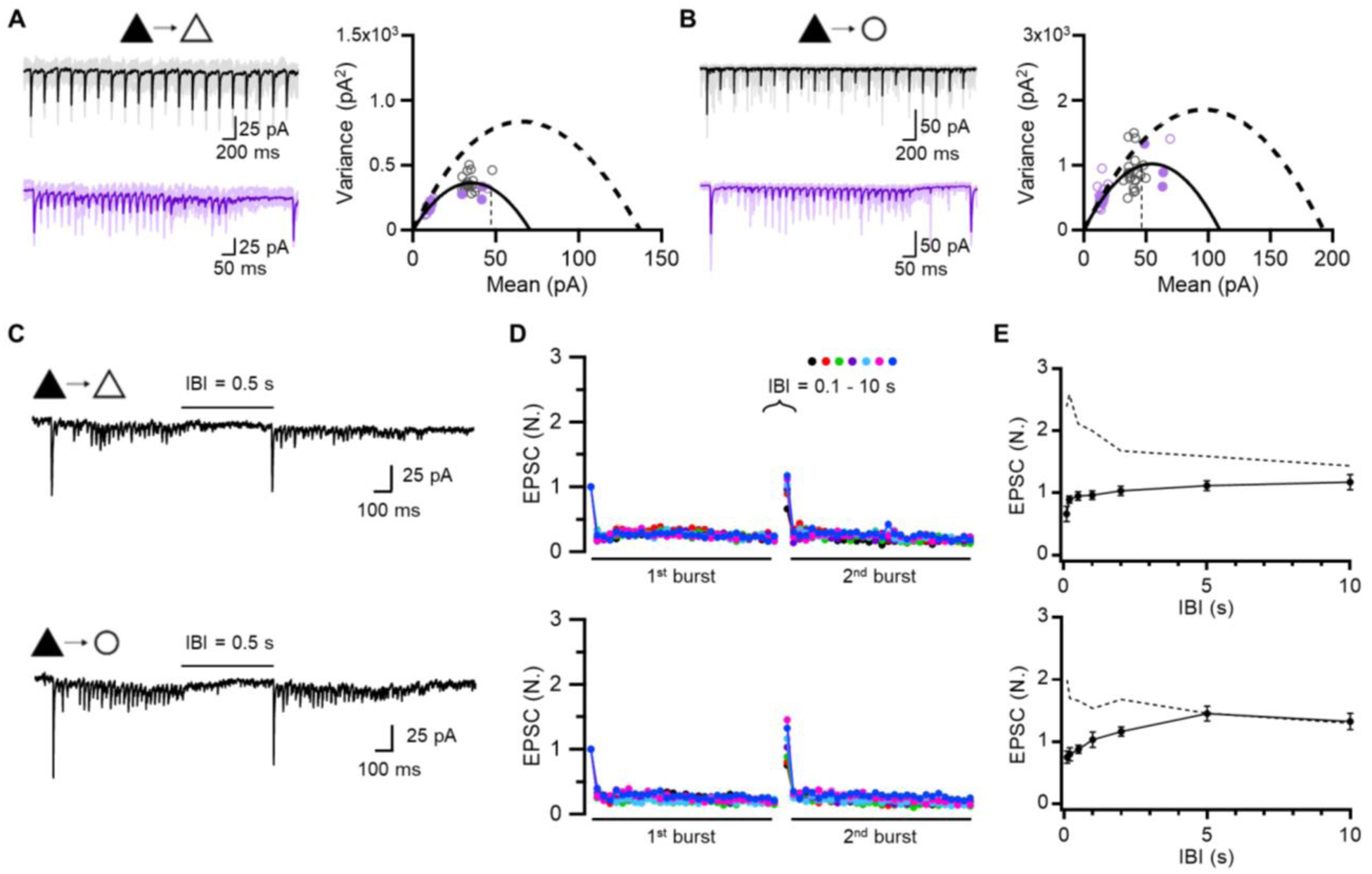
Syt7 KD synapses exhibit complementary changes in the number of release sites and their vesicle occupancy. **(A,B**) *Left*, Representative traces of EPSCs evoked by 5 and 40 Hz train stimulations (*black*, 5 Hz; *purple*, 40 Hz). *Right*, Variance-mean plots of EPSCs amplitude from averaged EPSCs recorded at PC-PC (*A*; n = 9, 15, 12, 10, 9) and PC-FSIN (*B*; n = 4, 14, 10, 9, 8) synapses in which presynaptic Syt7 has been knocked down. The data were fitted using MPFA and error bars are omitted for clarity. The mean 1^st^ EPSC amplitude of 5 Hz train (*vertical broken line*) was used for estimation of baseline p_occ_. Dashed parabolas indicate MPFA fits to variance-mean plot of WT synapses reproduced from Figure 3. (**C-E**) Recovery experiments at PC-PC (*top*) and PC-FSIN (*bottom*) synapses in which presynaptic Syt7 was knocked-down. (**C**) Representative EPSCs evoked by double 40 Hz train stimulations separated by 0.5 s. (**D**) Mean baseline-normalized amplitudes of EPSCs evoked by double 40 Hz trains at PC-PC (n = 12, 10, 9, 9, 10, 9, 8) and PC-FSIN (n = 10, 9, 10, 11, 8, 9, 10) synapses at different interburst intervals (IBIs). (**E**) Recovery time course. Baseline-normalized amplitudes of 1^st^ EPSC from the 2^nd^ burst were plotted as a function of various IBIs. *Dotted lines* indicate augmented EPSCs in the WT reproduced from Figure 3.

### TS vesicle recovery following depletion is accelerated by Syt7

Syt7 is recognized not only for its crucial role in facilitation, but also for mediating Ca^2+^-dependent replenishment of releasable vesicles (Liu et al., 2014; Tawfik et al., 2021). Given that facilitation at L2/3 excitatory synapses is driven by progressive overfilling (***Figure 2* *and* *Figure 3***), the loss of facilitation at Syt7 KD synapses in the present study implies that Syt7 plays a key role in the Ca^2+^-dependent acceleration of refilling and overfilling of the docking sites with TS vesicles. To test this hypothesis, we examined the recovery kinetics of EPSCs after TS vesicles were depleted by 3 pulses at 40 Hz in wild-type (WT) and Syt7 KD synapses (***Figure 7***). Strong PPD was observed not only in the first burst, but also in the second burst. This suggested that EPSCs in the second burst were mediated by TS vesicles, similar to those in the first burst (***Figure 7A, D***). Therefore, EPSC_1_ in the second burst can be interpreted as a release from TS vesicles recovering during a given IBI. The recovery time course of EPSC_1_ in the second burst indicated full recovery of TS vesicles within 100 ms in both excitatory synapse types in WT rats (***Figure 7C, F***). However, this recovery was remarkably slower in Syt7 KD synapses, requiring 5 s for full recovery, indicating that TS vesicle recovery was greatly accelerated by Syt7. Overall, synaptic facilitation and augmentation at local excitatory synapses in L2/3 can be ascribed to progressive activity-dependent increases in the occupancy of TS vesicles (***Figure 2* *and* *Figure 3***), in which Syt7 plays a key role.

**Figure 7.**
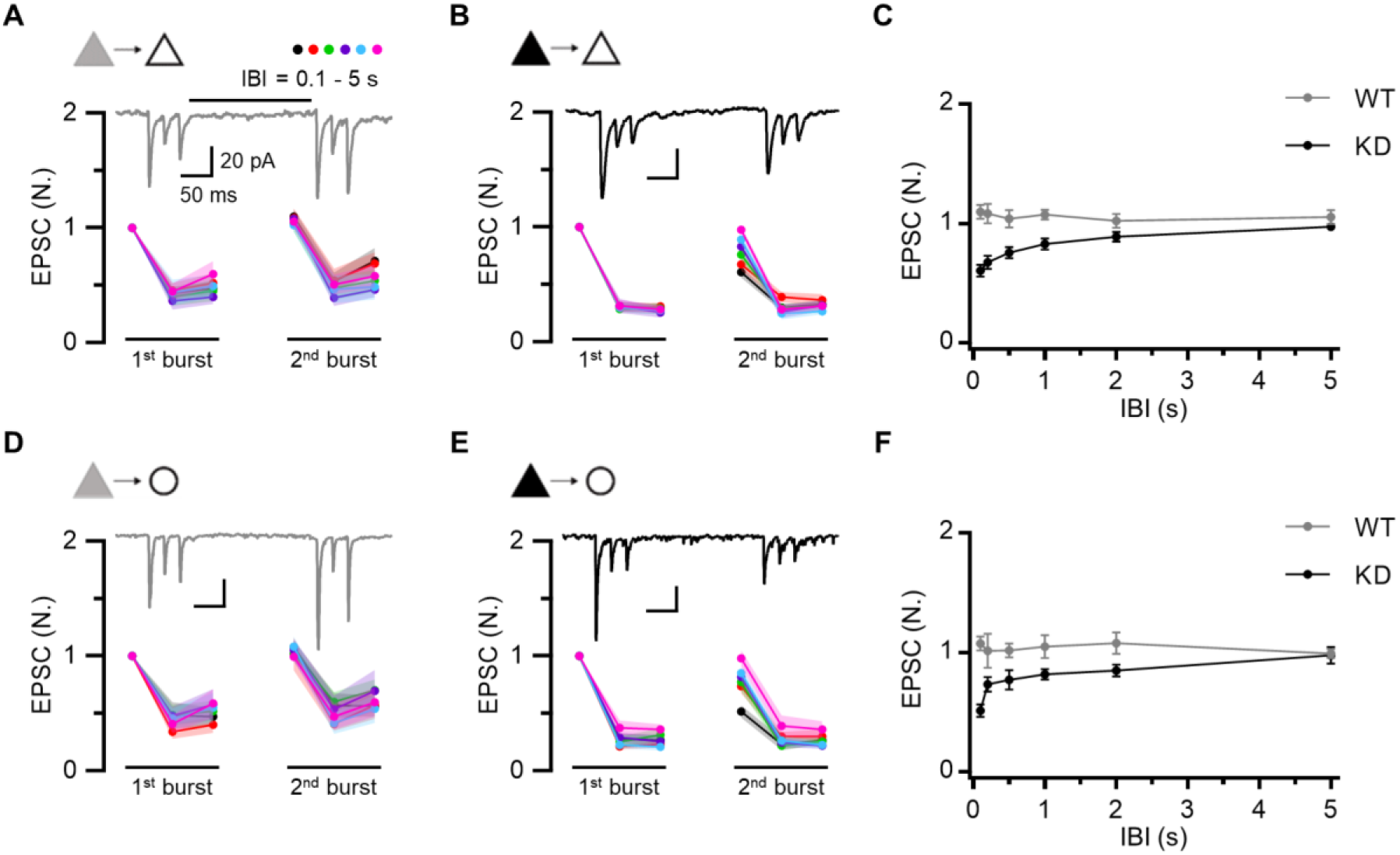
Recovery of TS vesicles following depletion is accelerated by Syt7. (**A-C**). Recovery experiments at PC-PC synapses in WT (*A*) and Syt7 KD (*B*). Mean baseline-normalized amplitudes of EPSCs evoked by two consecutive 3-pulse 40 Hz trains in WT (n = 16, 13, 11, 11, 11, 9 from short to long IBIs, respectively) and KD (n = 13, 11, 12, 8, 10, 11) synapses at different IBIs (0.1, 0.2, 0.5, 1, 2, 5 s). *Inset*, representative traces of EPSCs evoked by two consecutive 3-pulse 40 Hz train stimulations separated by 0.2 s. (**D-F**). Recovery experiments at PC-FSIN synapses in WT (*D*) and Syt7 KD (*E*). Mean baseline-normalized amplitudes of EPSCs evoked by two consecutive 3-pulse 40 Hz trains in WT (n = 9, 11, 11, 9, 10, 13) and KD (n = 9, 12, 9, 7, 10, 9) synapses at different IBIs. *Inset*, representative traces of EPSCs evoked by two consecutive 3-pulse 40 Hz train stimulations separated by 0.2 s. (**C, F**) Recovery time course. Baseline-normalized amplitudes of 1^st^ EPSC from the 2^nd^ burst were plotted as a function of various IBIs.

### Behavioral consequences of Syt7 KD in L2/3 PCs of the mPFC

We previously proposed that the disparity in STP between the PC-PC and PC-IN synapses results in activity-dependent changes in the ratio of synaptic weights at the PC-PC and PC-IN synapses (J_ee_/J_ie_). This may in turn have a profound influence on persistent activity in a recurrent network by biasing the E-I balance (Yoon et al., 2020). Syt7 KD not only eliminated facilitation, but also the activity-dependent increase in the J_ee_/J_ie_ ratio in the L2/3 network (***Figure 8—figure supplement 1***). This suggests the possibility that Syt7 contributes to the persistent activity during working memory tasks in the L2/3 recurrent network. Trace fear conditioning (tFC) requires associative learning between an auditory cue (conditioned stimulus, CS) and temporally separate aversive events (unconditioned stimulus, US). The formation of trace fear memory requires the prelimbic area of the mPFC and hippocampus. mPFC activity during the trace interval is thought to temporarily hold CS information, enabling the network to associate temporally discontinuous events (i.e., temporal associative learning) (Gilmartin et al., 2013). Synaptic facilitation and augmentation are implicated in temporary memory retention in recurrent networks (Mongillo et al., 2008; Mongillo et al., 2012).

Given that Syt7 is crucial for synaptic facilitation (***Figure 5***) and augmentation (***Figure 6***) in prelimbic L2/3 recurrent excitatory synapses, we tested whether trace fear memory formation was impaired in rats with depleted Syt7 transcripts, specifically in the L2/3 PCs of the mPFC. Syt7-targeted shRNA (shSyt7) or scrambled shRNA (Scr) constructs were transfected bilaterally into the L2/3 PCs of the mPFC using IUE (***Figure 8A***). Both the Scr-transfected control and shSyt7-transfected rats underwent tFC training (***Figure 8B***). Consistent with Gilmartin et al. (2013), in which prefrontal persistent firing was optogenetically inhibited, WT and KD animals exhibited similar freezing behavior during this acquisition phase. The following day, the rats were subjected to a tone test in which trace fear memory formation was assessed from the freezing response to a tone alone in a distinct context. Compared to the control group, the shSyt7-transfected group exhibited significantly lower levels of freezing behavior in response to auditory cues during the first four trials, suggesting an impairment in trace memory formation (***Figure 8C***).

**Figure 8.**
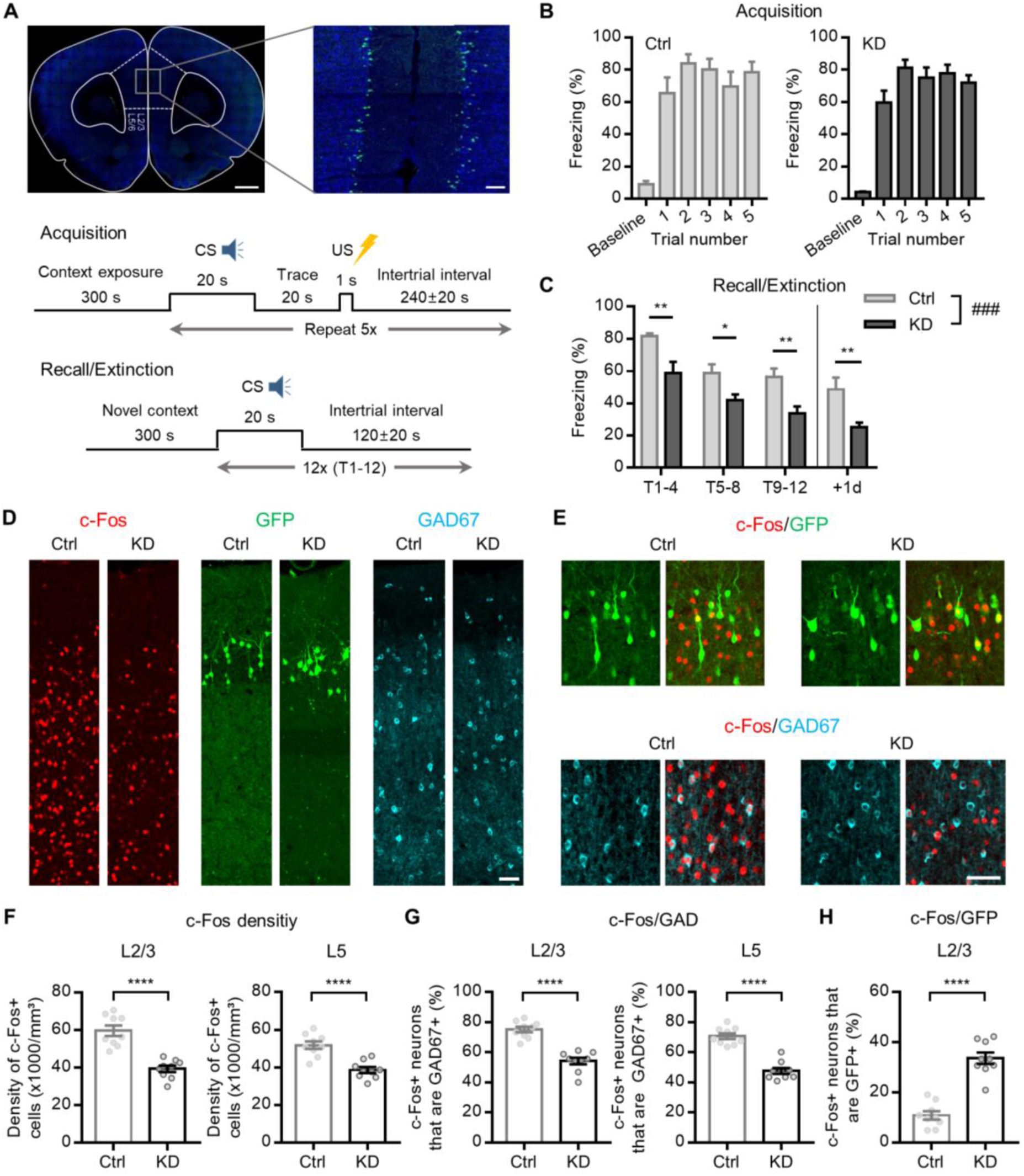
Behavioral effects of Syt7 deficiency in L2/3 PCs of the mPFC. (**A**) *Top*, Representative images showing bilateral expression of U6-GFP in L2/3 of PCs after IUE at E17.5. Scale bar: 1 mm, 100 μm. *Bottom*, Schematic of trace fear conditioning and extinction (tone test) protocol. (**B**) Freezing behavior of control (expressing scrambled shRNA, *Scr*) and Syt7 KD rats during acquisition of tFC. (**C**) Freezing ratio during tone tests on following days. Data are shown as average freezing during T1–T4, T5–T8, or T9–T12 (T, trials). The freezing on T1-T4 was significantly lower in KD rats suggesting that formation of trace memory was impaired in Syt7 KD rats (n = 10, 11 for Ctrl and KD, respectively; P = 0.0007, F(1, 19) = 16.43, two-way repeated measures ANOVA; P = 0.0064, 0.0211, 0.0064, 0.0064; Holm-Sidak test). (**D**) Representative images of c-Fos immunoreactivity in the prelimbic cortex of control or Syt7 KD rats 90 min after tFC acquisition. c-Fos (*left*, *red*, Cy5) and GAD67 (*right*, *cyan*, Cy3) were immunostained in the same brain slice expressing U6-GFP (*middle*; *green*). Scale bar, 50 μm. (**E**) Exemplar images of c-Fos positive neurons expressing GFP (*top*) or GAD67 (*bottom*) in control (*left*) or Syt7 KD rats (*right*). Scale bar, 50 μm. (**F-H**) Effects of Syt7 KD on c-Fos density (*F*) and percentage of c-Fos positive neurons co-labeled with GAD67 (*G*) or GFP (*H*) in L2/3 or L5 of prelimbic cortex (n = 9, 9 for Ctrl and KD, respectively). *Open symbols*, individual data. All statistical data are represented as mean ± S.E.M.; ****, P<0.0001; unpaired t-test.

The tone test was repeated the following day to test for extinction memory formation. Syt7 KD animals exhibited much less freezing to the CS (+1d) than did WT animals. Considering the Rescorla-Wagner model (Yau and McNally, 2023), our results suggest that the strength of CS-US associative memory was weaker in KD animals than in WT animals; thus, the acquisition of trace fear memory was impaired by Syt7 KD in the L2/3 PCs of the mPFC.

Next, we assessed the locomotor activity and anxiety levels of rats using the open field test (OFT) and elevated plus maze test (EPM). Scr controls and Syt7 KD rats showed similar exploratory patterns, displaying comparable total distance moved and time spent in the periphery of the OFT or the open/closed arms of the EPM (***Figure 8—figure supplement 2***). Previous studies have suggested that the acquisition of trace fear memory requires elevated trace activity in prelimbic PCs (Gilmartin et al., 2013). Given that c-Fos, an immediate early gene, is upregulated following patterned activities in neurons, its expression is considered a reliable indicator of recent neuronal activation (Fields et al., 1997; Guzowski et al., 2005). To investigate the effect of Syt7 on neuronal activity *in vivo* during tFC, animals were sacrificed at the time of peak c-Fos protein expression, approximately 90 min after completion of the behavioral task (Arime and Akiyama, 2017).

To elucidate the activation patterns of specific neuronal populations, we performed immunohistochemical co-labeling of c-Fos and GAD67 (a GABAergic interneuron marker) upon tFC acquisition. The density of c-Fos(+) cells and the percentage of c-Fos(+) cells among GAD67(+) cells in all prelimbic layers were significantly lower in Syt7 KD rats than in control rats (***Figure 8F-G***). Additionally, analysis of the proportion of active cells among transfected L2/3 PCs (c-Fos+/GFP+) revealed that Syt7 KD L2/3 PCs were significantly more active than Scr-transfected PCs (***Figure 8H***). This unexpected hyperactivity of Syt7 KD PCs may be attributed to the high reciprocal connectivity between PCs and FSINs in the neocortex (Holmgren et al., 2003; Otsuka and Kawaguchi, 2009), as PCs with short-term depression would transmit less input to FSINs, which in turn would exert less feedback inhibition on reciprocally connected PCs. Collectively, these findings show that the slow activation of vesicle refilling supports various forms of facilitation at excitatory synapses in PFC L2/3, which is essential for neuronal activity during temporal associative learning.

## Discussion

The present study characterized STP at local excitatory synapses in the L2/3 network of the rat mPFC at physiological extracellular [Ca^2+^]. These synapses displayed initial strong depression and then delayed facilitation at 40 Hz, while monotonous slowly developing facilitation was observed at lower frequencies. Our study suggests that such delayed facilitation after a brief depression results from a high p_v_ (***Figure 2* *and* *Figure 3* *and* *Figure 3—figure supplement 2***), along with Ca^2+^-dependent delayed acceleration of vesicle refilling and overfilling (***Figure 2* *and* *Figure 3***). The two-fold increase in EPSC during 20 pulse train stimulations under the condition of such a high p_v_ suggests a Ca^2+^-dependent increase in the vesicle replenishment rate during a train; this then leads to overfilling of release sites beyond their incomplete basal occupancy (***Figure 3A-B***). The high p_v_ and strong PPD did not result from artificial bouton stimulation, AMPAR desensitization, or any potential distortion caused by optogenetic methods (***Figure 1—figure supplements 4 and 5***). Additionally, similar to short-term facilitation, PTA could be explained by an increase in the TS vesicle occupancy (***Figure 3C-E***), consistent with previous studies showing post-tetanic increases in the p_occ_ and/or vesicle pool size at various types of synapses (Lee et al., 2010; Vandael et al., 2020; Tran et al., 2023; Silva et al., 2024). These findings suggest that release sites are partially occupied at rest and that a progressive overfilling of release sites with TS vesicles underlies facilitation and augmentation at intracortical L2/3 excitatory synapses.

### Definition of vesicular release probability in the present and previous studies

Previous studies for post-tetanic augmentation (Stevens and Wesseling, 1999; Garcia-Perez and Wesseling, 2008) observed invariance of the RRP size after tetanic stimulation. In these studies, the RRP size was estimated by hypertonic sucrose solution or as the sum of EPSCs evoked 20 Hz/60 pulses train (denoted as ‘RRP_hyper_’). Because reluctant vesicles can be quickly converted to TS vesicles (16/s) and are released during a train (Lee et al., 2012), it is likely that the RRP size measured by these methods encompasses both LS and TS vesicles. In contrast, we assert high p_v_ based on the observation of strong PPD, failure rates upon paired stimulations at ISI of 20 ms (***Figure 2* *and* *Figure 3—figure supplement 2***). Given that single AP-induced vesicular release occurs from TS vesicles but not from LS vesicles, p_v_ in the present study indicates the fusion probability of TS vesicles. From the same reasons, p_occ_ denotes the occupancy of release sites by TS vesicles. Note that our study does not provide direct clue whether release sites are occupied by LS vesicles that are not tapped by a single AP, although an increase in the LS vesicle number may accelerate the recovery of TS vesicles. As suggested in Neher (2024), even if the number of LS plus TS vesicles are kept constant, an increase in p_occ_ (occupancy by TS vesicles) would be interpreted as an increase in ‘vesicular release probability’ if it was measured based on RRP_hyper_.

By the way, it should be noted that increases in both p_v_ and p_occ_ may contribute to PTA. Post-tetanic potentiation (PTP) and phorbol esters had similar effects on baseline EPSCs (Taschenberger et al., 2016), and PDBu increased both the TS vesicle pool size (1.9-fold) and fusion probability of TS vesicles (1.3-fold) at the calyx synapses (Lin et al., 2025). Therefore, it is possible that PTP may increase both p_occ_ and p_v_ of TS vesicles, although it has not been yet directly tested. Moreover, acceleration of the latency in the hypertonicity-induced vesicle release raises a possibility for an increase in fusion probability of TS vesicles (Stevens and Wesseling, 1999; Garcia-Perez and Wesseling, 2008), although it is still controversial that the acceleration of the release latency represents a reduction in the activation energy barrier for vesicle fusion (Schotten et al., 2015).

### Progressive overfilling of docking sites

Conventional models for facilitation assume an increase in p_v_ after an AP of RRP vesicles, which had been in a low p_v_ or reluctant state at rest (Dittman et al., 2000; Turecek et al., 2016). This facilitation model is unlikely to explain our results, because TS vesicles are predominantly released not only at rest but also in the facilitated state during or after train stimulation, as evidenced by the strong PPD (ISI, 25 ms) during a 5-Hz train (***Figure 2D-E***) and at different intervals after a 40-Hz/30 pulse trains (***Figure 3D***). Moreover, the PPD (ISI, 25 ms) remained pronounced during the early phase of facilitation (***Figure 7***), arguing against the possibility of slowly increasing the p_v_ of reluctant vesicles. These results suggested that facilitation was mediated by an activity-dependent progressive increase in occupancy of TS vesicles (TS occupancy). Facilitation through an increase in TS occupancy has been termed frequency facilitation, differentiating it from paired pulse facilitation that relies on a residual calcium-dependent transient increase in the p_v_ of releasable vesicles (Neher, 2024).

Assuming an increase in TS occupancy, to attain a two-fold increase in EPSC (***Figure 1***), the baseline occupancy of the docking sites by TS vesicles should be low, as estimated from the V-M plot (c.a. 30% in ***Figure 3***). However, it is unclear whether the remaining docking sites that are not occupied by TS vesicles are truly vacant or are occupied by reluctant LS vesicles that are not released by the AP or AP trains. TS vesicles can be supplied by the transformation of LS to TS vesicles, both of which reside at the same docking site (LS/TS model; reviewed in Neher, 2024). Alternatively, TS vesicles can be supplied from a distinct replacement pool (RS/DS model; reviewed in Silva et al., 2021). Further studies, including ultrastructural analyses of active zones, are required to address this issue.

Given the crucial role of Syt7 in facilitation (***Figure 5***), the facilitation mechanisms are closely related to how Syt7 mediates synaptic facilitation. Based on the slow kinetics of Syt7, previous computational modeling studies have proposed that Syt7 may mediate short-term facilitation through an activity-dependent increase in p_v_ (Turecek et al., 2016; Jackman and Regehr, 2017; Turecek et al., 2017; Norman et al., 2023). The present study, however, found that both facilitation and augmentation were mediated by overfilling of the release sites (***Figure 2* *and* *Figure 3***). Moreover, Syt7-KD synapses displayed slower recovery of TS vesicles (***Figure 7***). These results suggest that Syt7 is essential for activity-dependent overfilling of docking sites with TS vesicles. Whereas the baseline EPSC was not altered (***Figure 1—figure supplement 3***), and complementary changes in the number of docking sites and their baseline occupancy were observed in Syt7 KD synapses (***Figure 6***). These results raise a possibility that Syt7 may play a role in providing additional vacant docking sites that can be overfilled during facilitation. It remains to be elucidated whether the decrease in the number of docking sites at Syt7 KD synapses is related to the subcellular localization of Syt7 to the plasma membrane of the active zone (Sugita et al., 2001; Vevea et al., 2021).

### STP model in light of known Ca^2+^ binding kinetics of Syt7

Syt7 acts as an upstream Ca^2+^ sensor that facilitates activity-dependent replenishment (Liu et al., 2014; Bacaj et al., 2015; Chen et al., 2017; Tawfik et al., 2021). The present study shows that EPSCs are mediated by TS vesicles both at rest and during activity-dependent facilitation. Releasable vesicles are homogeneous, and Syt7-dependent overfilling is responsible for the activity-dependent enhancement of EPSCs. Thus, we tested whether slowly developing facilitation could be replicated within the framework of a simple refilling model based on the known Ca^2+^ binding kinetics of Syt7 (Brandt et al., 2012). To this end, we made following assumptions: (1) a Ca^2+^-dependent increase in the forward refilling rate (k_1_) requires a full Ca^2+^-bound form of Syt7, to which multiple Ca^2+^ ions can bind (two or three Ca^2+^ ions on each C2A and/or C2B domain) (Brandt et al., 2012; Chon et al., 2015; Voleti et al., 2017); (2) sequential Ca^2+^ binding steps are synergistic through a cooperativity factor, *b*; (3) Syt7 senses AP-induced local [Ca^2+^] similar to that estimated at the calyx of Held (a Gaussian function with a full width at half maximum value of 0.2 ms and a peak of 40 μM) (Wang et al., 2008); and (4) residual [Ca^2+^] follows a mono-exponential function with an amplitude of 1 μM and decay time constant of 50 ms (Jackson and Redman, 2003).

Under these assumptions and the framework of the simple refilling model, slow and delayed facilitation could be replicated by modeling k_1_ as k_1_ = k_1,b_ + K_1,max_ × [Syt7: n Ca^2+^], where [Syt7: n Ca^2+^] represents the fraction of fully Ca^2+^-bound Syt7 (***Figure 9A-C***). Syt7 was slowly activated over the course of the stimulus train, suggesting that the cooperative binding of multiple Ca^2+^ ions progressively increased the putative active form of Syt7 because of its slow membrane binding/unbinding kinetics. Nevertheless, the late phase of facilitation at high frequencies was predicted to be higher in the simulation than in the experiment. As expected, based on the notion that sustained release during HFS is limited by site clearance in the active zone (***Figure 4C***), the model/data discrepancy became more pronounced at higher frequencies. Incorporating endocytic terms into the model may mitigate late-phase discrepancies. Moreover, removal of the catalytic function of Syt7 in the model effectively accounted for the complete loss of facilitation observed in Syt7 KD (***Figure 9D***).

**Figure 9.**
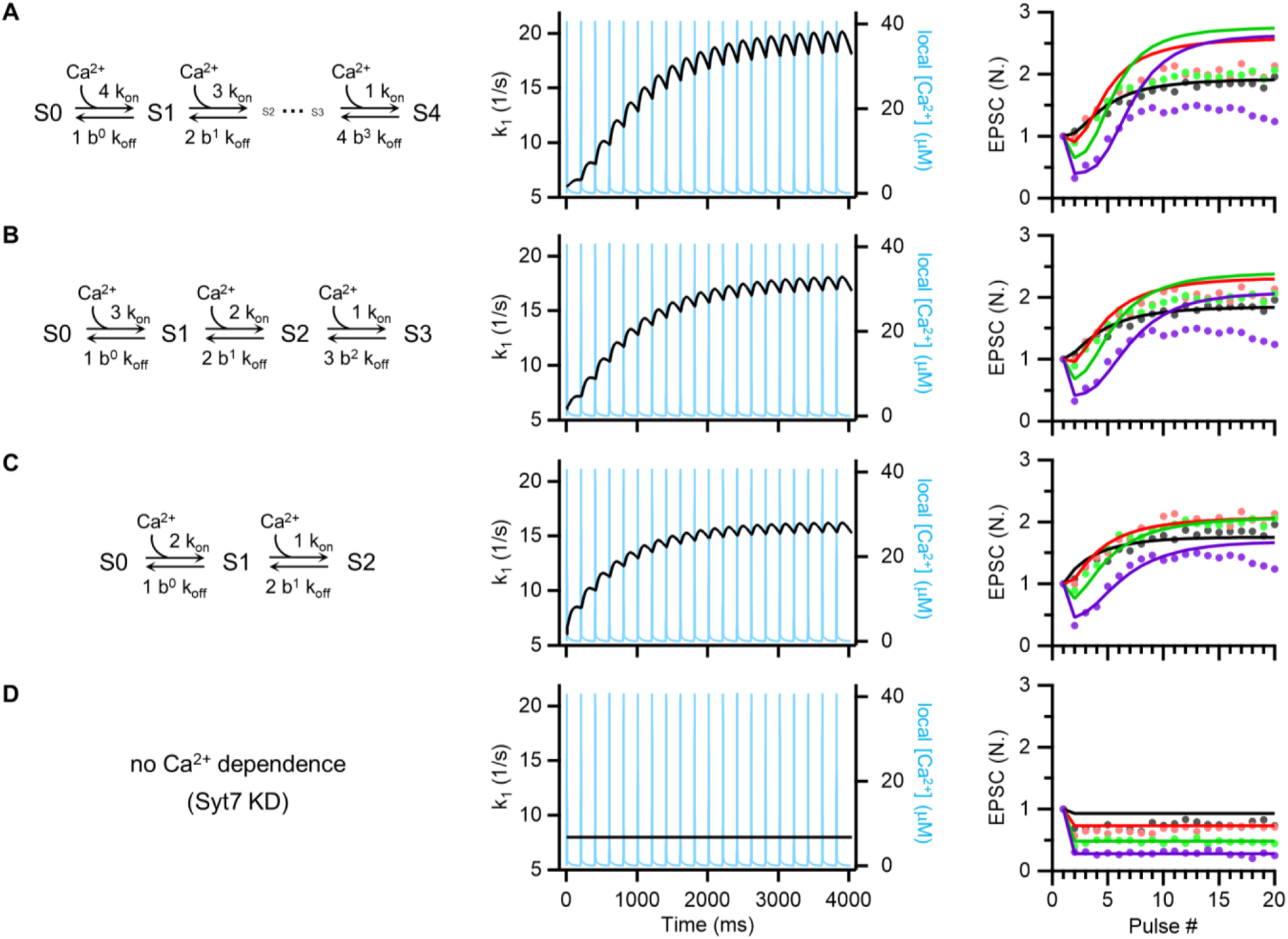
STP model in light of known Ca^2+^ binding kinetics of Syt7. (**A-D**) *Left*, Schematic of allosteric calcium binding to Syt7. The number of Ca^2+^ bound to Syt7 was denoted as # in ‘S#’ in the reaction scheme. k_on_ = 7/μM/s, k_off_ = 10/s. *Middle*, Simulated changes of k_1_ (*black*) in response to 5 Hz train of Ca^2+^ transients (*light blue traces*). The priming step of the simple refilling model was assumed to be catalyzed by full Ca^2+^-bound form of Syt7. Accordingly, Ca^2+^-dependent increase in k_1_ was calculated as K_1,max_ multiplied by a fraction of full Ca^2+^-bound form of Syt7. We assumed that local [Ca^2+^]_i_(t) follows a Gaussian function: (1/σ√2π) exp[-(t-t_p_)^2^/ 2σ^2^], in which t_p_ = 0.25 ms and σ = 0.085 ms. *Right*, Fits of the Syt7 model to the STP data. To fit this model to the STP data, K_1,max_ was set to 300/s (*A*), 220/s (*B*), and 180/s (*C*). Cooperativity factor (*b*) was set to 0.35 (*A*), 0.2 (*B*), and 0.05 (*C*). k_1_ is set to be constant for Syt7 KD (*D*). Same frequency color codes were used as in Figure 1.

### Conventional STF models involving cumulative increase in p_v_ of reluctant vesicles

Conventional models for facilitation assume a post-AP residual Ca^2+^-dependent step increase in p_v_ of RRP (Dittman et al., 2000) or reluctant vesicles (Turecek et al., 2016). Given that p_v_ of TS vesicles is close to one in the present study, an increase in p_v_ of TS vesicles cannot account for facilitation. The possibility for activity-dependent increase in fusion probability of LS vesicles (denoted as p_v,LS_) should be considered in two ways depending on whether LS and TS vesicles reside in distinct pools or in the same pool. Notably, strong PPD at short ISI implies that p_v,LS_ is near zero at the resting state. Whereas LS vesicles do not contribute to baseline transmission, short-term facilitation (STF) may be mediated by cumulative increase in p_v_ of LS vesicles that reside in a distinct pool. Because the increase in p_v,LS_ during facilitation recruits new release sites (increase in N), the variance of EPSCs should become larger as stimulation frequency increases, resulting in upward deviation from a parabola in the V-M plane, as shown in recent studies (Valera et al., 2012; Kobbersmed et al., 2020). This prediction is not compatible with our results of V-M analysis (***Figure 3***), showing that EPSCs during STF fell on the same parabola regardless of stimulation frequencies. Therefore, it is unlikely that an increase in fusion probability of reluctant vesicles residing in a distinct release pool mediates STF in the present study. For the latter case, in which LS and TS vesicles occupy in the same release sites, it is hard to distinguish a step increase in fusion probability of LS vesicles from a conversion of LS vesicles to TS. Nevertheless, our results do not support the possibility for a gradual increase in p_v,LS_ that occurs in parallel with STF. Strong PPD, indicative of high p_v_, was consistently found not only in the baseline (***Figure 2* *and* *Figure 2—figure supplement 1***) but also during post-tetanic augmentation phase (***Figure 3D***) and even during the early development of facilitation (***Figure 2D-E* *and* *Figure 7***), arguing against the gradual increase in p_v,LS_. One may argue that STF may be mediated by a drastic step increase of p_v,LS_ from zero to one, but it is not distinguishable from conversion of LS to TS vesicles.

### Possible mechanisms underlying the depression-facilitation sequence

Cooperative binding of multiple Ca^2+^ ions to Syt7 predicts delayed activation and a progressive increase in vesicle refilling during train stimulations, resulting in delayed facilitation. Combining this with a high p_v_ could replicate not only the slow facilitation at 5 Hz, but also the unique depression-facilitation sequence observed at 40 Hz (***Figure 9***). As simulated by Pulido and Marty (2018), delayed facilitation can be reproduced by a high p_v_ in combination with a low baseline occupancy of replacement sites and subsequent Ca^2+^-dependent refilling. However, whether this model is also able to replicate slow facilitation at low frequencies remains to be tested. An alternative, but not exclusive, possibility for delayed facilitation is that the presynaptic global Ca^2+^ itself may undergo a slow buildup during a train due to the saturation of endogenous Ca^2+^-binding proteins.

### Facilitation requires not only Syt7, but also activation of PLC-DAG signaling

To gain insight into the specific molecular mechanisms underlying the vesicle-loading processes, we examined the effects of pharmacological inhibitors targeting PLC and DAG (***Figure 4***). Both a PLC inhibitor and a DAG analog reduced facilitation and augmentation. This suggests that the progressive overfilling of docking sites with TS vesicles is supported by the PLC-DAG signaling pathway and probably downstream Munc13 (Rhee et al., 2002; Lou et al., 2008). Phorbol esters have been recently shown to increase the TS fraction of docked vesicles by enhancing vesicle priming without affecting Ca^2+^ transients or p_v_ (Taschenberger et al., 2016; Aldahabi et al., 2022; Papantoniou et al., 2023; Aldahabi et al., 2024). In line with these findings, signaling involving PLC and DAG may support Ca^2+^-dependent overfilling of TS vesicles by enhancing molecular priming processes.

### Roles of facilitation and augmentation in working memory and related tasks

Previous theoretical studies have proposed that working memory in a recurrent network can be maintained in the form of augmentation at synapses (Mongillo et al., 2008) and/or persistent activity of neurons in a memory ensemble (Mongillo et al., 2012). The latter can be supported by short-term facilitation of recurrent excitatory synapses or disparity in STP between excitatory and inhibitory synapses (Mongillo et al., 2012; Yoon et al., 2020). However, whether these short-term enhancements in synaptic efficacy contribute to working memory-related behaviors has not been examined. tFC is a temporal associative learning task requiring temporary retention of CS information during a trace period. The mPFC and hippocampus are crucial for tFC, and tFC is interfered with by a load of working memory and attention distraction, implying that the tFC shares the same neural substrate with other working memory tasks (Raybuck and Lattal, 2014). In the present study, tFC in WT and Syt7 KD animals revealed that Syt7 deficiency in the L2/3 PCs of the mPFC impaired the acquisition of trace fear memory and reduced c-Fos expression associated with tFC training (***Figure 8***).

The acquisition of trace fear memory was impaired by inhibition of persistent activity in mPFC during trace period (Gilmartin et al., 2013). The similar deficit observed in Syt7 KD animals is consistent with the hypothesis that STF provides bi-stable ensemble activity in a recurrent network (Mongillo et al., 2012). Nevertheless, alternative mechanisms may be responsible for the behavioral deficit. Not only recurrent network but also long-range loop between the mPFC and the mediodorsal (MD) thalamus play a critical role in maintaining persistent activity within the mPFC especially for a delay period longer than 10 s (Bolkan et al., 2017). Prefrontal L2/3 is heavily innervated by MD thalamus, and L2/3-PCs subsequently relay signals to L5 cortico-thalamic (CT) neurons (Collins et al., 2018). Given that L2/3 is an essential component of the PFC-thalamic loop, loss of STF at recurrent synapses between L2/3 PCs may lead to insufficient L2/3 inputs to L5 CT neurons and failure in the reverberant PFC-MD thalamic feedback loop. Therefore, not only L2/3 recurrent network but also its output to downstream network should be considered as a possible network mechanism underlying behavioral deficit caused by Syt7 KD L2/3.

In summary, activity-dependent synaptic facilitation and augmentation at prelimbic L2/3 recurrent excitatory synapses can be attributed to overfilling of release sites with TS vesicles. Given that the lack of short-term facilitation in L2/3 PCs impairs trace fear memory acquisition, temporary maintenance of CS information during trace period may be supported by an increase in the TS occupancy in presynaptic terminals of CS-representing neuronal ensembles.

## Materials and Methods

### Animals/Subjects

All experiments were carried out on Sprague-Dawley rats of either sex at p28-43. Rats were maintained in temperature-controlled rooms on a constant 12 h light/12 h dark cycle and were given access to water and food ad libitum. All animal procedures were approved and performed in accordance with the Institutional Animal Care and Use Committee at Seoul National University (SNU200522-1).

### In utero electroporation (IUE)

Pregnant Sprague Dawley rats on embryonic day (E) 17.5 were deeply anesthetized with 5% (v/v) isoflurane for the duration of surgery. The uterine horns were exposed by laparotomy and wet with warm sterile phosphate-buffered saline (PBS). The plasmids (0.5-2 µg/µl) together with the 0.01% fast green dye were injected into either the left or the right lateral ventricle or bilateral ventricles of embryos through a thin glass capillary (Narishige PC-10). The embryo’s head was carefully held between the forceps-shaped electrodes of 10 mm diameter (CUY65010; NEPAGENE, Japan). Electroporation pulses (50 V for 50 ms) were delivered five times at intervals of 150 ms using a square-wave generator (ECM830; BEX, United States). The uterine horns were placed back into the abdominal cavity and abdominal muscles and skin were sutured. After the surgery, the animals were recovered under the infrared lamp. For knockdown experiments in slice electrophysiology, the shRNA constructs were co-electroporated with the pCAG-oChIEF-tdTomato, because of the consistent co-expression of the two constructs in a significant proportion of the transfected cells in vivo (Bony et al., 2013). Pups with robust fluorescence signals in the prefrontal cortex of both hemispheres were used for behavior experiments.

### Preparation of vector constructs

For an optic stimulation, pCAG-oChIEF-tdTomato was transfected by IUE. Short hairpin RNA (shRNA) against Syt7 sequence (GATCTACCTGTCCTGGAAGAG) was expressed under the control of an U6 promoter of pAAV-U6-GFP vector (Cell Biolabs, #VPK-413). This shRNA was shown to effectively knockdown Syt7 in cell cultures and in vivo by previous reports from more than one research group (Bacaj et al., 2013; Li et al., 2017). Scrambled control sequence (TCGCATAGCGTATGCCGTT) was designed by shuffling the recognition region sequences and a BLAST analysis confirmed that the control shRNA sequence had no target gene. For rescue experiments, the cDNA for rat Syt7 (NM_021659) rendered resistant to the shRNA were inserted in the KD vector and their expression was driven by the CAG promoter; the vector also contained a self-cleaving P2A peptide sequence followed by EGFP, which ensures visualization of transfected cells. All the constructs above were verified by DNA sequencing. Plasmid DNA was purified and concentrated under endotoxin free conditions (Qiagen EndoFree Maxi Kit).

### Slice preparation

Acute brain slices were prepared by using a VT1200S Vibratome (Leica Microsystems) after isoflurane anesthesia and decapitation. Coronal sections of the mPFC (300 μm in thickness) were cut in ice-cold artificial cerebrospinal fluid (aCSF) composed of (in mM) 125 NaCl, 3.2 KCl, 25 NaHCO_3_, 1.25 NaH_2_PO_4_, 20 D-glucose, 2 NaPyr, 1 MgCl_2_, and 1.3 CaCl_2_ (Yoon et al., 2020), bubbled with 95% O_2_ and CO_2_ (pH = 7.3; 310 mOsm). Slices were allowed to recover for a minimum of 30 minutes at 36 °C, and subsequently maintained at room temperature until recording. Composition of aCSF was identical throughout the preparation, incubation, and recording periods unless otherwise stated. For the recordings, the mPFC slices were continuously perfused with oxygenated aCSF at a flow rate of 1.5 ml/min, and the temperature was maintained at 31–32 °C using an in-line temperature controller (Warner Instruments).

### Electrophysiology

Whole-cell patch-clamp recordings were performed from pyramidal cells and fast-spiking interneurons in layer 2/3 of the prelimbic mPFC. Neuronal subtypes were initially selected based on morphology under visual guidance by using an upright microscope equipped with infrared differential interference contrast (IR-DIC) optics (BX51WI, Olympus), and were further identified by their electrophysiological properties (see ***Figure 1—figure supplement 1***; van Aerde and Feldmeyer, 2015; Tremblay et al., 2016). To analyze these intrinsic properties in current-clamp experiments, we measured the following parameters: (1) resting membrane potential (RMP), (2) input resistance (R_in_), (3) sag ratio, and (4) the F-I curve, which plots firing frequencies (F) against the amplitude of injected currents (I). Both cell types exhibited a low sag ratio below 0.1, consistent with findings from other studies (van Aerde and Feldmeyer, 2015; Yoon et al., 2020). PCs and FSINs were easily distinguished by their unique firing frequency, Rin, spike adaptation, RMP and morphological characteristics including the shape of soma and the thickness of apical dendrites. For experiments involving oChIEF-expressing neurons in WT or Syt7 KD rats, whole-cell recordings were performed from the fluorescently labeled pyramidal neurons in layer 2/3 of the prelimbic cortex using suitable filter sets. Patch pipettes (tip resistance of 2–3 MΩ) were pulled from borosilicate glass and filled with (in mM) 130 K-gluconate, 8 KCl, 10 HEPES, 0.2 EGTA, 20 Na_2_Pcr, 0.3 Na_2_GTP, 4 MgATP. Intracellular solutions were adjusted to pH 7.25 and 300–310 mOsm. Neurons were voltage clamped at −78 mV and series resistance (Rs) were continuously monitored during the EPSCs recordings. Experiments were discarded before analysis if the Rs changed by 20% or larger deviation of baseline value. Measurements were not corrected for Rs compensation, bridge balance, or liquid junction potential. Electrophysiological data were low-pass-filtered at 1 kHz (Bessel) and acquired at 20 kHz using a Multiclamp 700B amplifier paired with Digidata 1440A digitizer and Clampex 10.2 software (Molecular Devices).

### Synaptic stimulation

IUE resulted in sparse expression of channelrhodopsin (10-20%) (***Figure 1A***). Exploiting the sparse expression, we employed a minimal optical stimulation method to activate a single or very few excitatory synapses in layer 2/3. The optic minimal stimulation was performed by our published protocol with minor modifications (Yoon et al., 2020). This approach aimed to minimize the risk of synaptic contamination, which could arise from engaging extra terminals that were initially inactive (depolarized below threshold) but became active by temporal summation during high-frequency trains. It also helped prevent the buildup of depolarization at the synaptic terminals, which might influence the probability of release, as noted by (Jackman et al., 2014). To achieve this, we employed a collimated digital micromirror device (DMD)-coupled LED (Polygon400; Mightex Systems) to confine 470 nm blue light to a small area with a radius of approximately 3–8 μm (typically 3–4 μm), as measured at the focal plane of a 60× water immersion objective [numerical aperture (NA) = 1.0; LUMPlanFL, Olympus] (***Figure 1—figure supplement 3A***). To find the minimal stimulation area, the radius of illumination area was increased from 2 μm by 50% at each step (***Figure 1—figure supplement 3A***), and selected the smallest illumination area that elicited EPSCs. Setting the duration of illumination between 3 and 5 ms (typically 5 ms), photostimulation onto oChIEF-expressing cells reliably induced action potentials (APs) during 600-pulse trains (see ***Figure 1—figure supplement 2***).

STP at various stimulus frequencies (5, 10, 20, and 40 Hz) was examined using 20-pulse trains. Trials of train stimulation were repeated with a prolonged interval of 30 s to avoid change in baseline properties. We observed no systematic changes in the amplitudes of the first EPSC of each train, nor did we find significant differences in the paired-pulse ratio (PPR) across stimulation trials in the same cell. For each stimulus frequency, approximately ten to twenty consecutive traces were collected and then averaged across trials to yield a mean EPSC trace. For recovery experiments, two consecutive trains of 3 or 30 stimuli at 40 Hz, with varying time intervals (0.1-10 s), were applied every 10 s or 60 s, respectively, at both excitatory synapses. For 600-stimulus trains at 10Hz, the trial was delivered only once for each cell to avoid possible after-effects caused by prolonged synaptic stimulation (Hirsch and Crepel, 1990). Prior to the stimulus trains, the EPSCs were normalized to the mean amplitude of the baseline. For the baseline measurements, paired pulses with various ISIs (20-200 ms) were delivered every 3 s (1/3 Hz) at least 20 times.

For all recordings, picrotoxin (PTX, 10 μM) was included in the recording solution to isolate glutamatergic responses. mPFC PC terminals were entirely glutamatergic as the application of 6-cyano-7-nitroquinoxaline-2,3-dione (CNQX, 20 μM), an AMPA receptor antagonist, completely abolished the monosynaptic events evoked by photostimulation. A low concentration of PTX was employed to minimize the relieving effects on spike suppression caused by blockade of GABAa receptors. We confirmed that no synaptic response was elicited by electrical stimulation in the presence of both 10 μM PTX and 20 μM CNQX.

In ***Figure 1—figure supplement 4A***, 1 µM TTX (1 mM stock solution) and 0.1 mM 4-AP (500 mM stock solution) were added to the bath solution to block action potentials and restore presynaptic glutamate release, respectively (***Figure 1—figure supplement 4***). In ***Figure 1—figure supplement 5A***, 50 µM of cyclothiazide, a positive allosteric modulator of the AMPA receptors, was included to minimize rapid desensitization of the postsynaptic receptors. In ***Figure 1—figure supplement 5B***, 0.5 mM kynurenate was added to bath solution to prevent possible saturation of AMPA receptors, especially during maximal synaptic facilitation. For pharmacological manipulations of synaptic vesicle dynamics (***Figure 4***), drugs related to vesicle supply including U73122, OAG, LatB, EGTA-AM, and dynasore at indicated concentrations were added to the recording solution during baseline measurements or PTA/recovery experiments (IBI = 0.5 s). Chemicals were from Sigma-Aldrich, Merck, or Tocris.

### Measurements of quantal size

Experiments for measuring asynchronous release were performed as previously described (Yoon et al., 2020). In brief, we replaced extracellular Ca^2+^ entirely with 2 mM Sr^2+^ to induce asynchronous release. A single stimulus was delivered at 3-second intervals over 100 repetitions, and asynchronous release events were detected during a 100-ms window starting 25 ms after stimulus onset for each trial. The same procedure was also executed under sham conditions (0% light intensity) to serve as a control, thus mitigating contamination from nonspecific spontaneous events and potential errors associated with in the detection algorithm. q was determined as the mean value from the first Gaussian function after fitting the EPSC distribution with either a single or double Gaussian function (***Figure 3—figure supplement 1***).

### Estimation of Ca^2+^ kinetics at axonal boutons

Cytosolic [Ca^2+^] at single axonal boutons was estimated by employing the two-dye, dual-excitation wavelength technique as described by (Oheim et al., 1998). This method involves measuring the fluorescence ratio of a Ca^2+^-sensitive dye to that of a Ca^2+^-insensitive dye to correct for dye concentrations. We used Fluo-5F at concentrations of 150, 250, or 500 μM (dissociation constant, K_d_ = 2.3 μM) as the calcium indicator dye and Alexa Flour 555 (50 μM) as the reference dye. The fluorophores were from Invitrogen. Both dyes were co-loaded into presynaptic terminals of layer 2/3 pyramidal cells in mPFC through whole-cell patch clamp. Before data acquisition, the dyes were allowed to diffuse to terminal boutons for at least 30 minutes after the patch break-in, a duration sufficient for the equilibrium of dye concentrations between the soma and its axonal terminals.

Calcium imaging was performed by sequentially exciting Fluo-5F and AF555 at wavelengths of 473 and 559 nm, respectively, using a confocal laser-scanning microscope (FV1200; Olympus, Japan) equipped with a 60× water immersion objective (NA, 0.9; LUMPlanFl; Olympus, Tokyo, Japan). Once a single axonal bouton was identified, lines for scanning were drawn across boutons, oriented perpendicular to the axon (***Figure 2—figure supplement 2A-B***). After baseline fluorescence was measured, line scanning was proceeded while a single or train APs were induced by applying current pulses. Upon stimulation, green fluorescence but not red one increased along the vertical array of lines across the bouton (***Figure 2—figure supplement 2B-C***). Measurements were obtained under current-clamp conditions, with the resting potential regularly monitored. The scanning speed was adjusted to 600-700 Hz by controlling the line length. Line scan for measuring fluorescent transients in a bouton was repeated at least 3 times, and then averaged to enhance the signal-to-noise ratio. To mitigate photobleaching, minimal intensity of the laser power and maximal pinhole size were used. Emitted light was filtered through 490/590 or 575/675 bandpass filters before detection by photomultiplier tubes for green or red fluorescence, respectively.

Fluorescence measurements were converted to [Ca^2+^]_i_ using the ratio of background-subtracted fluorescence (F = F_0_ - F_b_) from two dyes. The fluorescence ratio (R = F_green_/F_red_) in data traces were converted to [Ca^2+^]_i_ using the following equation:

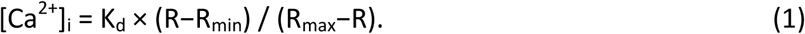

Calibration parameters were obtained using an in-cell calibration protocol (Helmchen, 2011), wherein the minimum (R_min_) and maximum ratio (R_max_) values were established through the use of intracellular solutions containing 10 mM EGTA or 10 mM [Ca^2+^], respectively.

When Ca^2+^ is extruded with a rate constant, γ, after an AP-induced short pulse of Ca^2+^ influx to a single compartment, [Ca^2+^]_i_ initially rises to A_Ca_ and then undergoes a mono-exponential decay with a time constant (τ_Ca_). Under this framework, A_Ca_ and τ_Ca_ depend on Ca^2+^ binding ratios of endo- and exogenous Ca^2+^ buffers (denoted as κ_S_ and κ_B_, respectively) as following equations (Neher, 1995; Kim et al., 2003):

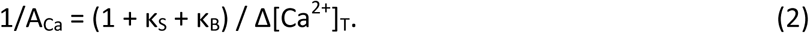

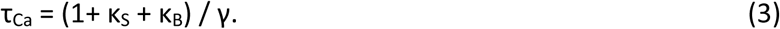

, where Δ[Ca^2+^]_T_ is an increment in total [Ca^2+^] (free plus bound) elicited by an AP. Ca^2+^ binding ratio indicates the ratio of concentration changes in the Ca^2+^-buffer complex relative to free [Ca^2+^] change, and linearized κ_B_ is calculated from the dye concentration and the change from [Ca^2+^]_1_ to [Ca^2+^]_2_ as follows:

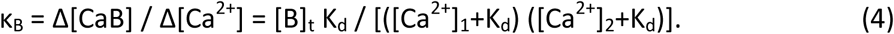

The total dye concentration, [B]_t_, was taken as [Fluo-5F] in the patch pipette. With κ_B_ determined from Equation 4 and τ_Ca_ from the decay of a Ca^2+^ transient, κ_S_ can be estimated by plotting τ versus κ_B_ and applying a linear fit to the plot, which gives the x-intercept as -(1 + κ_S_). Once determined, κ_S_ can be used to calculate Δ[Ca^2+^]_T_.

### Optimization of k_1_ and p_v_ from failure rates

Let n_0_ = the number of docked SVs before arrival of a first AP; n_1_ = the number of docked SVs remaining after a first AP (i.e. n_0_ - n_1_ vesicles are released); n_2_ = the number of docked SVs just before arrival of a 2^nd^ AP (ISI = 20 ms; ***Figure 3—figure supplement 2***). If the number of release sites (N), forward and reverse rates for vesicle docking (k_1_ and b_1_, respectively), mean release site occupancy (p_occ_) and vesicular fusion probability upon an AP (p_v_) are given, we can predict P(n_0_, n_1_, n_2_), the probability for [n_0_, n_1_, n_2_], as following steps.

1) Probability for a synapse harbouring n_0_ docked SVs just before arrival of a first AP is calculated as:

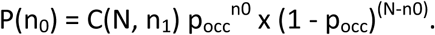

2) Probability that n_1_ vesicles remain after a first AP when the initial number of docked SVs is n_0_ [P(n_1_|n_0_)] is calculated as:

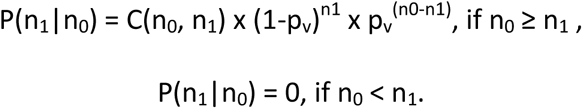

3) When forward and backward rate constants are k_1_ and b_1_, respectively, probability that n_2_ vesicles are docked just before arrival of a 2^nd^ AP when n_1_ SVs remain after a 1^st^ AP (Δt = 20 ms) [P(n_2_ | n_1_)] is predicted as follows: Assuming that docking and undocking follow a Poisson process,

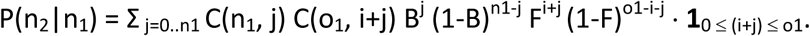

where B = 1 – exp(-b_1_ Δt); F = 1 – exp(-k_1_ Δt); i = n_2_ – n_1_; o_1_ = N-n_1_.

4) Combining probabilities calculated from above three steps, we can calculate P(n_0_, n_1_, n_2_):

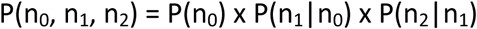

5) Once P(n_0_, n_1_, n_2_) is given, we can predict probability for double failure (P_00_) and probability for the 1^st^ success and the 2^nd^ failure (P_10_) as:

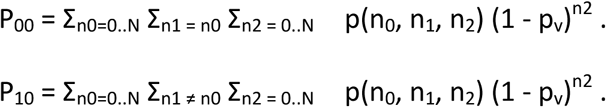

6) Probability for the 1^st^ failure and the 2^nd^ success (P_01_) and probability for double success (P_11_) were calculated as:

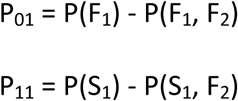

in which P(F_1_) and P(S_1_) are probability for the 1^st^ failure and the 1^st^ success, respectively, and these are calculated as:

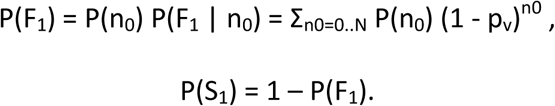

To optimize k_1_ and p_v_, we used the Matlab routine, *fmincon*, in which the sum of squared errors (= observed value minus predicted value) was set as a cost function to be minimized. Lower and upper bounds for k_1_ and p_v_ were set as [4, 8] and [0.8, 1], respectively. Release probability (p_r_) was calculated from the observed 1^st^ failure rate, P(F_1_), as

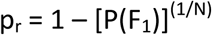

N was assumed to be six as shown in ***Figure 3***. Other parameters (b_1_, and p_occ_) were set according to following relationships:

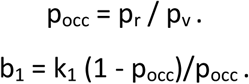

### Calculation of probability for double failure

Double failure probability, P_00_, can be more easily predicted. By the Bayes rule,

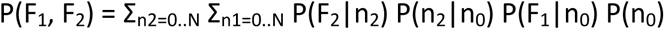

where F_1_ and F_2_ means 1^st^ and 2^nd^ failure, because n_0_ = n_1_ after the 1^st^ failure. P(F_1_) and N are observed values. For calculation in ***Figure 3—figure supplement 2***, k_1_ and p_v_ were set free variables. Other parameters were set under the following constrains: P(F_1_) = (1 – p_r_)^N^, p_r_ = p_v_ p_occ_, and p_occ_ = k_1_ / (k_1_ + b_1_).

### Behavioral tests

#### Trace fear conditioning

For trace fear conditioning (tFC) (Gilmartin et al., 2013), individual rats were placed in the training chamber (30.5 x 25.4 x 30.5 cm; Coulbourn Instruments) consisting of metal grid floor connected to an electrical stimulator (H10-11R-TC-SF, Coulbourn Instruments) for delivery of a footshock. After a 5 min habituation period, rats received five pairings of a 20 s tone conditioned stimulus (CS, 2.5 kHz, 75 dB), followed by empty 20s trace period and a unconditioned stimulus (US; 1 s, 0.7 mA footshock) with a pseudorandom inter-trial interval (ITI) of 240 ± 20 s. During this training session, rats learned to associate the auditory CS with the shock US. The next day, the rats were tested for memory of CS-US trace association (called tone test). The acquisition of fear learning, often indicated by the extent of freezing to the CS, was assessed in a novel chamber (a white hexagonal enclosure) in a dimly lit room. During this tone (or retrieval) test, rats were first allowed to explore the new context for 5 min (habituation), followed by twelve 20 s CS presentations with a variable ITI of 120 ± 20 s. This entire session was repeated on the next day for assessment of extinction learning. Identities of the chambers were also determined with the presence of distinct odors; the training and testing chambers were cleaned with 70% ethanol and 1% acetic acid respectively before and after each session. For all experimental sessions, the activity of animals in the chambers was recorded at 30 frames per second using the EthoVision XT (Noldus Information Technology, Wageningen, Netherlands), and stored as a video file. The freezing behavior, defined as behavioral immobility other than respiratory movement, was assessed using an open-source video analysis pipeline ezTrack (Pennington et al., 2019). The percentage of freezing was calculated as a total duration of freezing divided by the total duration of observation.

#### Elevated Plus Maze

Elevated plus maze (EPM) test was performed using a plus-shaped apparatus elevated 60 cm above the ground. The maze consisted of two open arms (50 x 10 x 0.5 cm (H)), two closed arms (50 x 10 x 40 cm (H)) and a center area (10 x 10 cm). Rats were individually placed on the center area, facing an open arm, and the path of the animal was recorded with a video camera for 5 min. We analyzed the total distance traveled for 5 min and time spent in each arm. Animals were tracked offline using the open-source tracking system ezTrack (Pennington et al., 2019).

#### Open field test

Open field exploration test (OFT) was performed in an open field apparatus in a dimly lit room. The open field consisted of a white plastic board (1.20 m in diameter) surrounded by white plastic walls (50 cm in height). Individual rats were placed in a center zone (0.6 m in diameter), and allowed to freely explore the apparatus for 10 min. Time spent in the center zone and total distance traveled were analyzed for initial 5 min. Animals were tracked offline using the open-source tracking system ezTrack (Pennington et al., 2019).

### Histology and immunohistochemistry

At the end of the behavioral experiments, rats were anesthetized using isoflurane inhalation and transcardially perfused with PBS followed by 4% paraformaldehyde (PFA, T&I, Korea). Brains were extracted and post-fixed in 4% PFA solution for 24 h. Perfused brains were sliced at 100 μm for estimation of GFP expression or 50 μm for immunohistochemistry. For analysis of c-Fos expression, rats were returned to their home cages immediately after the training session and perfused approximately 90 minutes later (Arime and Akiyama, 2017). Brains were then cryopreserved, frozen, and sectioned coronally. Immunofluorescence staining was conducted on free-floating slices using a rabbit monoclonal anti c-fos (1:1000, Cell Signaling Technology, cat# 2250s) and mouse monoclonal anti-GAD67 (1:500, Millipore, cat# mab5406). Fluorescent secondary antibodies included Cy5-conjugated goat anti-rabbit IgG (1:1000, Abcam, cat# ab97077) and Cy3-conjugated goat anti-mouse IgG (1:1000, Abcam, cat# ab97035). Brain slices were permeabilized and blocked with PBS containing 0.5% Triton X-100 and 5% normal goat serum (NGS) for 1h at room temperature. Slices were incubated overnight at 4 °C with primary antibodies, followed by 4 h incubation with secondary antibodies at room temperature. Slices were then mounted with Antifading Mounting medium (Abcam, cat# ab104139). Confocal images were acquired using a Leica TCS SP8 microscope set to the same laser intensity and acquisition parameters to compare the immunosignals in sections from Scr and Syt7 KD groups. For cell counting and colocalization, fluorescent protein-positive cells were automatically quantified with spot-detection algorithms in Imaris 9.5 (Oxford instruments).

### Data analysis

Data were analyzed and presented using ClampFit (Molecular Devices), Igor Pro (Wavemetrics), MATLAB (Mathworks), and Prism (GraphPad). Data points and error bars represent the mean ± standard error of the mean. All statistical analyses were conducted using two-tailed comparisons. Sample sizes and statistical tests for each group are detailed in the figure legends.

### Numerical integration of the simple refilling model

This model posits reversible docking of vesicles at finite release sites (N_max_) with forward (k_1_) and reverse rate constants (b_1_) (Hosoi et al., 2007; Neher and Sakaba, 2008).

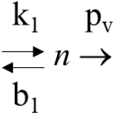

According to this scheme, d*n*/dt = k_1_ *u* - b_1_ *n* - p_v_ *n* δ(t - t_AP_), where *u* is the number of unoccupied sites (= N_max_ - *n*); t_AP_ is the timing of AP firing; and δ is the Dirac delta function. The p_v_ was assumed to be unity (***Figure 3—figure supplement 2***). The sum of basal k_1_ (= k_1,b_) and b_1_ was be estimated as 23/s from the dependence PPR on ISIs (***Figure 3—figure supplement 2Ab***). From the baseline occupancy of 0.3 and p_occ_ = k_1,b_ / (k_1,b_ + b_1_), we could estimate baseline k_1_ and b_1_ as 6.9/s, and 16.1/s, respectively. Ca^2+^_-_ and Syt7-dependent increase of k_1_ was modeled as k_1_ = k_1,b_ + K_1,max_ [Syt7: n Ca^2+^], where [Syt7: n Ca^2+^] represents the fraction of full Ca^2+^-bound Syt7. For deterministic simulations, we time-integrated a set of differential equations for each model using Euler methods with a time step of 1 ms. All calculations were performed on the platform of Matlab (R2022b, Mathworks, USA).

### Material availability statement

All data generated or analyzed during this study are included in the manuscript and supporting files. Source data files are provided in this paper. Matlab codes for the analyses and simulation are presented in this paper.

## Acknowledgements

We thank Dr. E. Neher and Dr. A. Marty for critical reading of this manuscript and invaluable comments.

## Author contributions

All experiments were carried out at cell physiology lab under Y.K and S.H.L’s supervision. Conception or design of the work: J.S. and S.H.L. Acquisition, analysis or interpretation of data for the work: J.S, Y.K. and L.S.H. Drafting the work or revising it critically for important intellectual content: J.S., S.Y.L, Y.K. and L.S.H.

## Supplementary Figures

**Figure 1—figure supplement 1.**
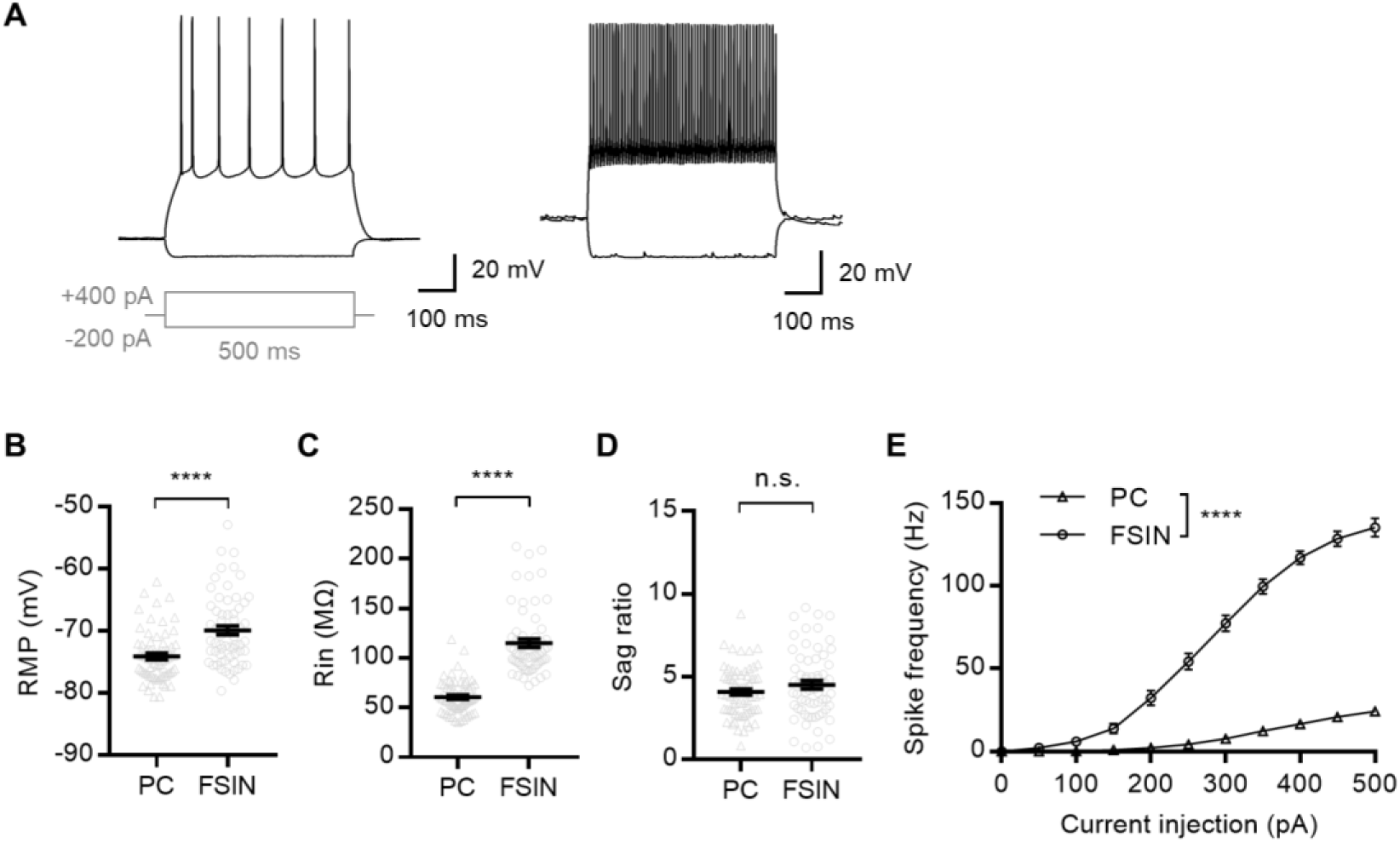
Intrinsic membrane properties of mPFC neurons. (**A**) Neuronal subtypes in mPFC layer 2/3. Cells were characterized based on their electrophysiological properties (n = 66, 64 for PC and FSIN, respectively). Representative voltage responses to current injection (-200 or +400 pA) of pyramidal cells (*left*) and fast-spiking interneurons (*right*). (**B-D**) Mean resting membrane potential (*B*), input resistance (*C*), and Sag ratio (*D*) measured in layer 2/3 PCs and FSINs. *Gray symbols*, individual data. (**E**) Current-spike relationship in each cell type. All statistical data are represented as mean ± S.E.M.; n.s. = not significant; ****, P<0.0001; unpaired t-test or two-way ANOVA.

**Figure 1—figure supplement 2.**
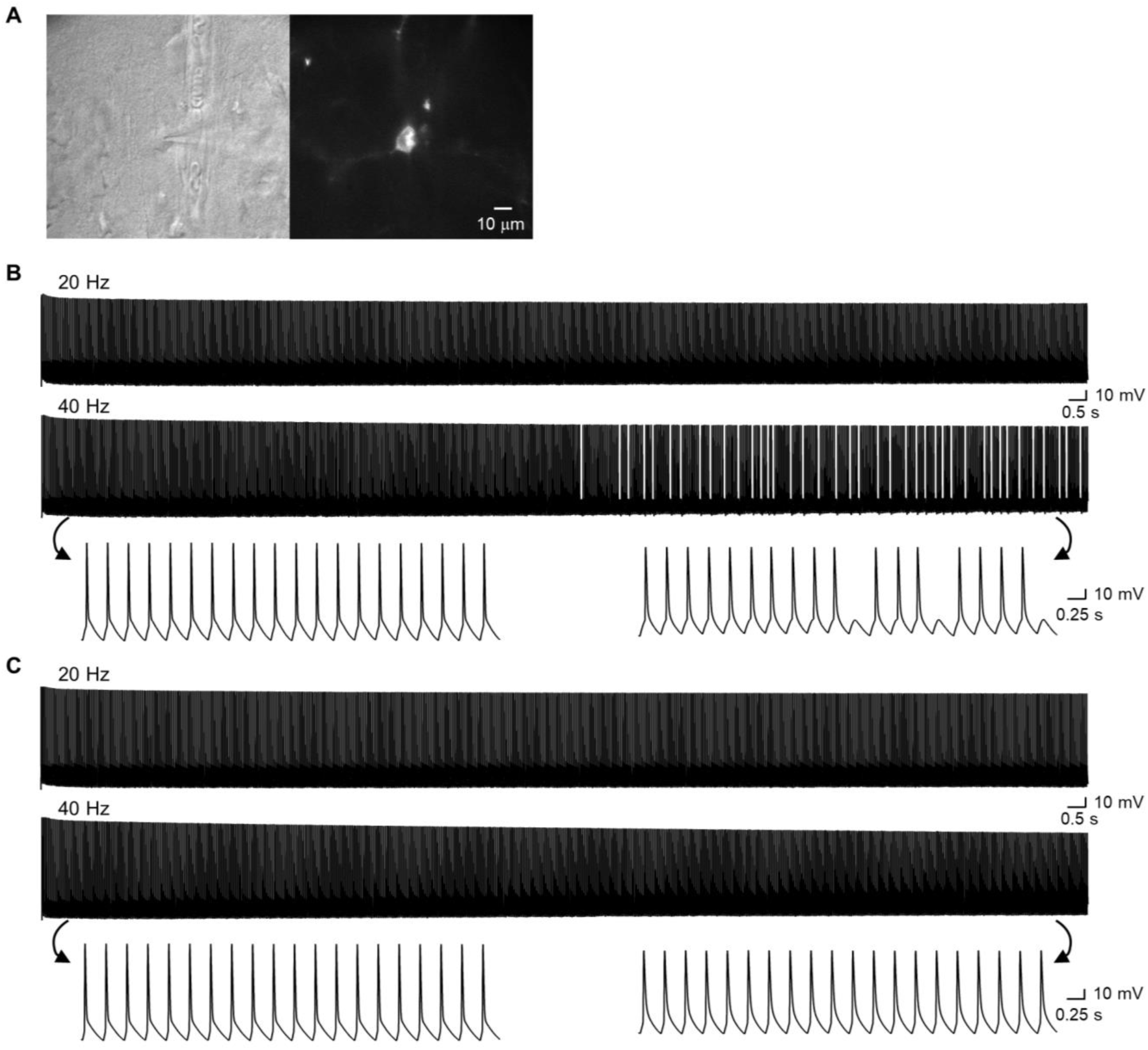
Test for consistency of light stimulation in oChIEF-expressing L2/3 pyramidal cells. (**A**) Representative images of an oChIEF-expressing neuron. Whole-cell recordings were performed from the fluorescently labeled neurons in layer 2/3 of the prelimbic cortex. *Left*, IR-DIC image (60×) of the pyramidal cell under whole-cell patch clamp. *Right*, fluorescence image of td-Tomato in the same cell excited at 530 nm wavelength. (**B**) Current-clamp recordings of action potentials evoked by 600 light pulse trains from an oChIEF-expressing L2/3 cell. For somatic stimulation, we used the typical light stimulation conditions used in STP experiments (5 ms in duration and 4 μm in radius; see *Methods*). Uniform action potentials were produced with high fidelity, albeit not without compromise towards the end of the train in the case of 40 Hz stimulation. Note that these results are similar to previously reported values from neocortical neurons (Lin et al., 2009; Yoon et al., 2020). (**C**) Same experiments as (*B*) but in Syt7 KD pyramidal cells that express oChIEF in L2/3 of the mPFC.

**Figure 1—figure supplement 3.**
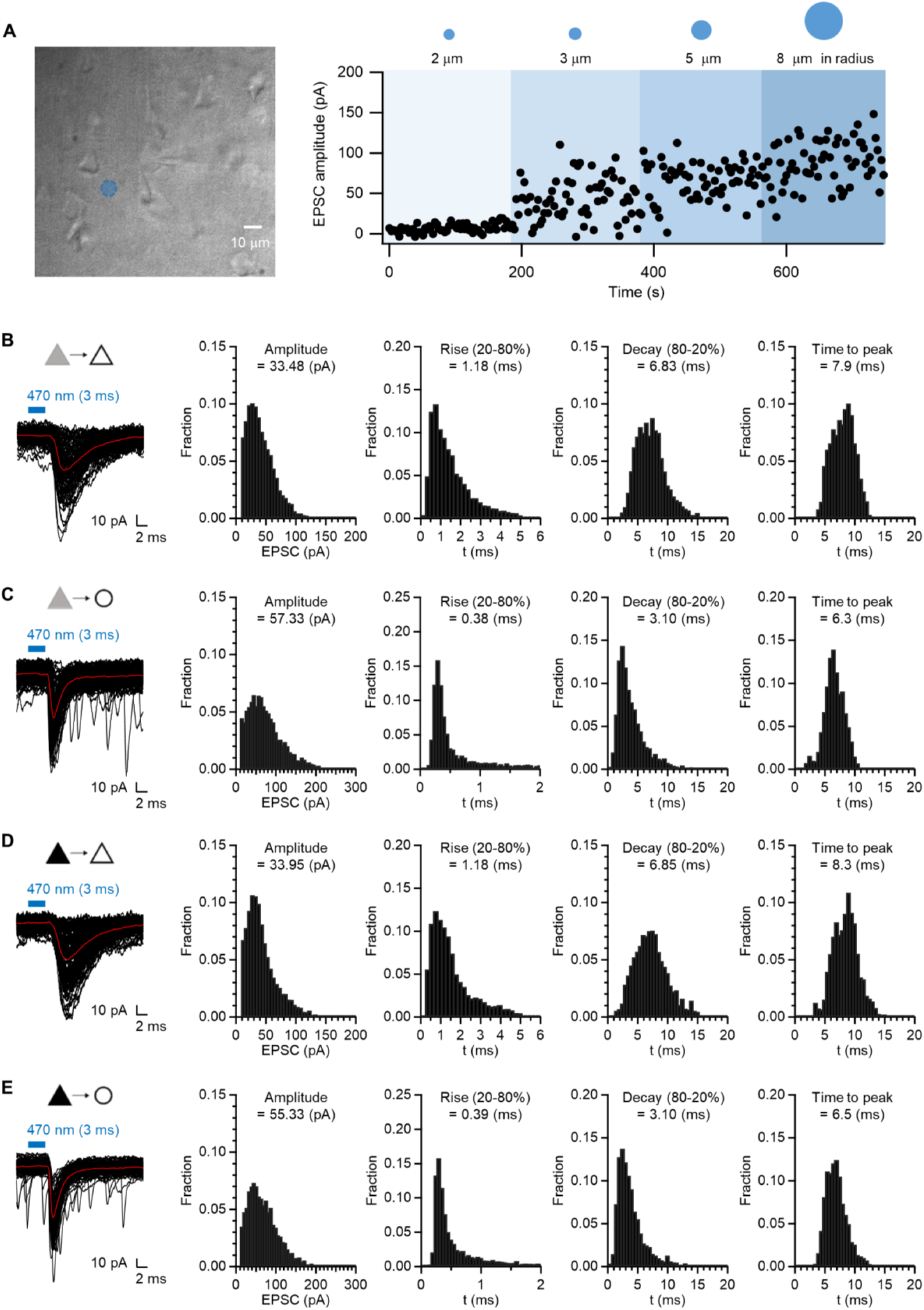
Properties of EPSCs evoked by minimal stimulation. (**A**) Optical minimal stimulation. A collimated DMD-coupled LED was used to confine the area of excitation to a small region near the soma. To stimulate a minimal number of excitatory axon fibers, the radius of illumination area was increased from 2 μm by 50% at each step, and selected the smallest illumination area that elicits EPSCs (Yoon et al., 2020) before starting each recording session. The illumination area was in the range of 3 to 8 μm in radius, when measured at the focal plane (typically 3–4 μm) of a 60× water immersion objective (*blue circle* in the photograph). (**B-E**) Properties of minimally stimulated baseline EPSCs (failure-excluded). Medians of frequency distributions are presented. Decay times below 1 ms were excluded from analysis. From left to right, representative traces (100 pulses), EPSC amplitude, rise time (20-80%), decay time (80-20%), and time to peak (from stimulus onset) from WT PC-PC (*B*, n = 81), WT PC-FSIN (*C*, n = 77), Syt7 KD PC-PC (*D*, n = 72), and Syt7 KD PC-FSIN (*E*, n = 78) synapses.

**Figure 1—figure supplement 4.**
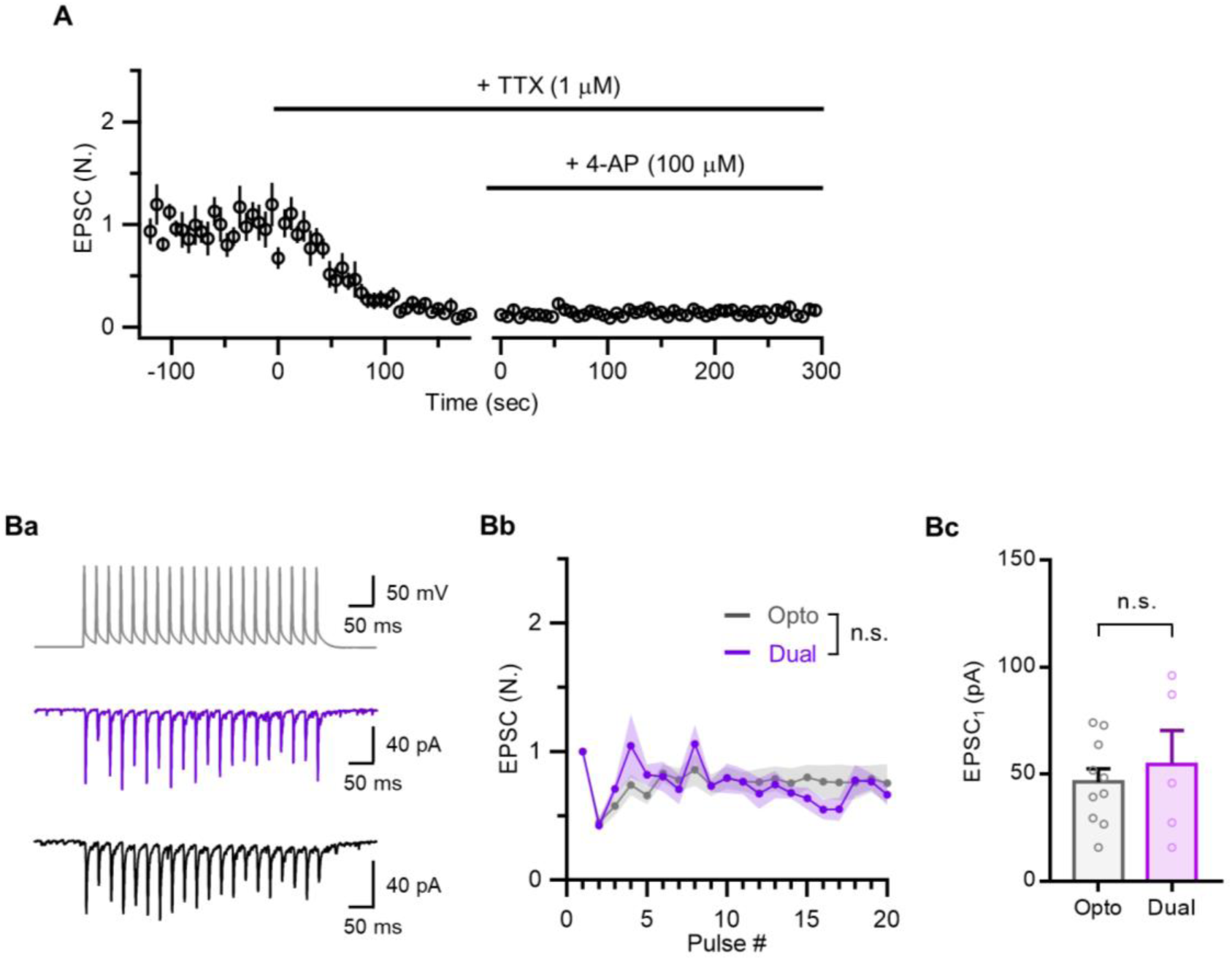
Validation of STP of optically evoked EPSCs. (**A**) The high p_v_ at PC-PC synapses may result from direct photo-stimulation of axon boutons. To test if optically evoked EPSCs depend on APs, the effect of 1 µM tetrodotoxin (TTX) was tested. Applying 1 μM tetrodotoxin (TTX) and additionally 0.1 mM 4-aminopyridine, a D-type K channel blocker abolished optically evoked EPSCs (n = 8). Even when repolarization was hindered by subsequent addition of 100 µM 4-aminopyridine (4-AP), a potassium channel blocker, EPSCs were not rescued. These findings suggest that the light-evoked responses are dependent on APs of presynaptic axon fibers rather than photo-induced depolarization of presynaptic boutons (Little and Carter, 2013). (**B**) Comparison of STP evoked by optical stimulation with that by dual patch techniques. To test validity of STP of photostimulation-evoked EPSCs, we compared STP of unitary EPSCs evoked by dual whole-cell patch at PC-FSIN synapses. Both of EPSC1 and STP at 40 Hz recorded by using paired whole-cell recordings were not significantly different from those evoked by photo- stimulation, supporting the validity of our optogenetic methods. (**Ba**) Representative EPSC traces evoked by 20-pulse 40 Hz trains stimulation at pyramidal cell to fast spiking interneuron (PC-FSIN) synapses. A train of APs in a presynaptic PC (*top*), evoked EPSCs in a FSIN by dual whole-cell patch clamp (*middle*) or by optical stimulation (*bottom*) under same conditions. (**Bb**) Mean values for baseline-normalized amplitudes of EPSCs evoked by 40 Hz train at PC-FSIN synapses. *Opto*, optical stimulation (n = 10); *Dual*, dual patch techniques (n = 5). STP evoked by the two techniques were not significantly different (two-way repeated measures ANOVA, F (1, 13) < 0.01, P = 0.955). (**Bc**) Mean values for the first EPSC in different stimulation methods (P = 0.5724, unpaired t-test). *Open* c*ircles*, individual data. All statistical data are represented as mean ± S.E.M.

**Figure 1—figure supplement 5.**
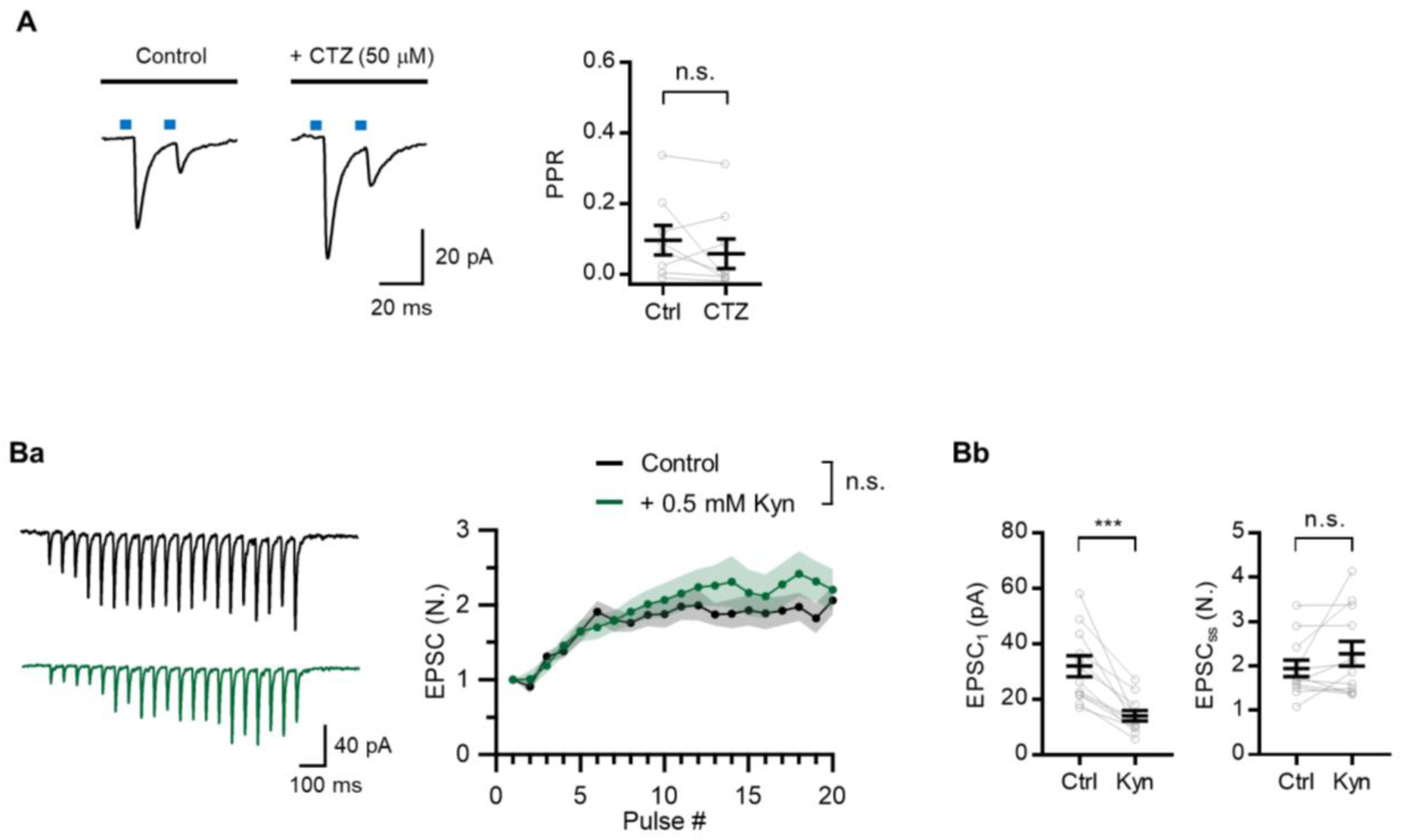
Test for AMPAR desensitization and saturation. (**A**) Representative traces of EPSCs evoked by paired-pulse photostmulation at 20 ms ISI (*left*) and mean values for paired pulse ratio (PPR, *right*) before and after applying 50 μM cyclothiazide (CTZ, n = 8), an AMPA receptor (AMPAR) desensitization inhibitor. CTZ induced no significant change in PPR arguing against the possibility that strong PPD is caused by AMPAR desensitization. *Gray symbols*, individual data. (**B**) It has been shown that multi-vesicular release occurs at recurrent excitatory synapses in neocortical L2/3 (Holler et al., 2021). Whereas a single vesicle release is insufficient to saturate postsynaptic AMPARs, an increase in the number of vesicular release during facilitation may saturate AMPAR resulting in underestimation of facilitation. To test this possibility, facilitation at 20 Hz was measured in the presence of 0.5 mM kynurenic acid (Kyn), a fast competitive antagonist (Diamond and Jahr, 1997; Wadiche and Jahr, 2001). Fast blockade of AMPA receptors did not have significant effects on short-term facilitation at excitatory synapses in the mPFC L2/3, suggesting that AMPAR saturation may not be significant during facilitation. (**Ba**) Representative EPSC traces (*left*) and average of baseline-normalized EPSCs (*right*) evoked by 40 Hz 20-pulse trains in control and in the presence of 0.5 mM kynurenic acid (Kyn). Although facilitation was slightly increased by Kyn, it was not statistically significant (n = 12; two-way repeated measures ANOVA, F (1, 22) = 0.575, P = 0.456). The slightly higher facilitation may result from reduced blocking efficiency of Kyn upon increased glutamate concentration in the synaptic cleft (Tong and Jahr, 1994; Meyer et al., 2001). (**Bb**) Mean values for the baseline EPSC (*left*) and the steady-state EPSC (*right*) before and after applying 0.5 mM kynurenate as shown in *Ba*. The baseline EPSC was reduced by Kyn to 43.5% of control. *Gray symbols*, individual data. All statistical data are represented as mean ± S.E.M.; n.s. = not significant; ***, P<0.001; paired t-test.

**Figure 2—figure supplement 1.**
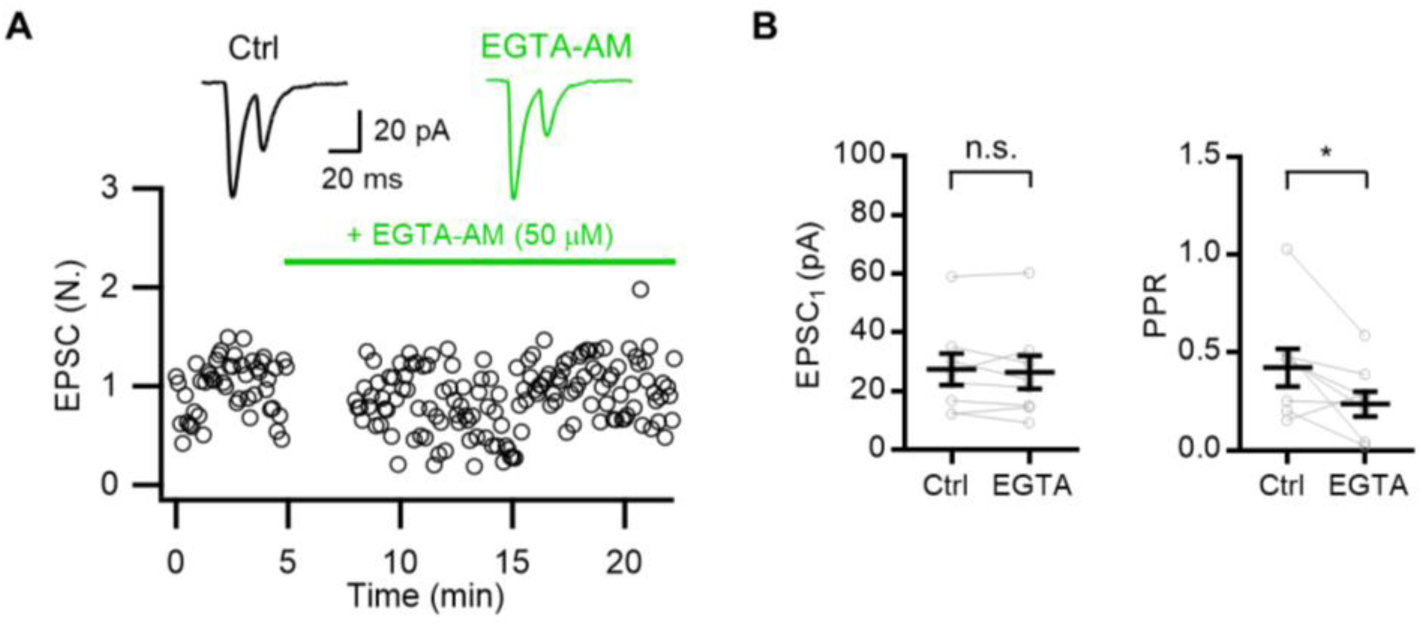
Effects of EGTA-AM on synaptic transmission. (A) Representative traces (*top*) and time courses of normalized amplitudes of the first EPSCs (*bottom*) during paired-pulse photostimulation at 20 ms ISI in control and in the presence of 50 μM EGTA-AM (n = 8). The green horizontal bar indicates application of 50 μM EGTA-AM. (B) Mean values for the first EPSC (*left*) and PPR (*right*) before and after EGTA-AM application. *Gray symbols*, individual data. All statistical data are represented as mean ± S.E.M.; n.s. = not significant; *, P<0.05; paired t-test.

**Figure 2—figure supplement 2.**
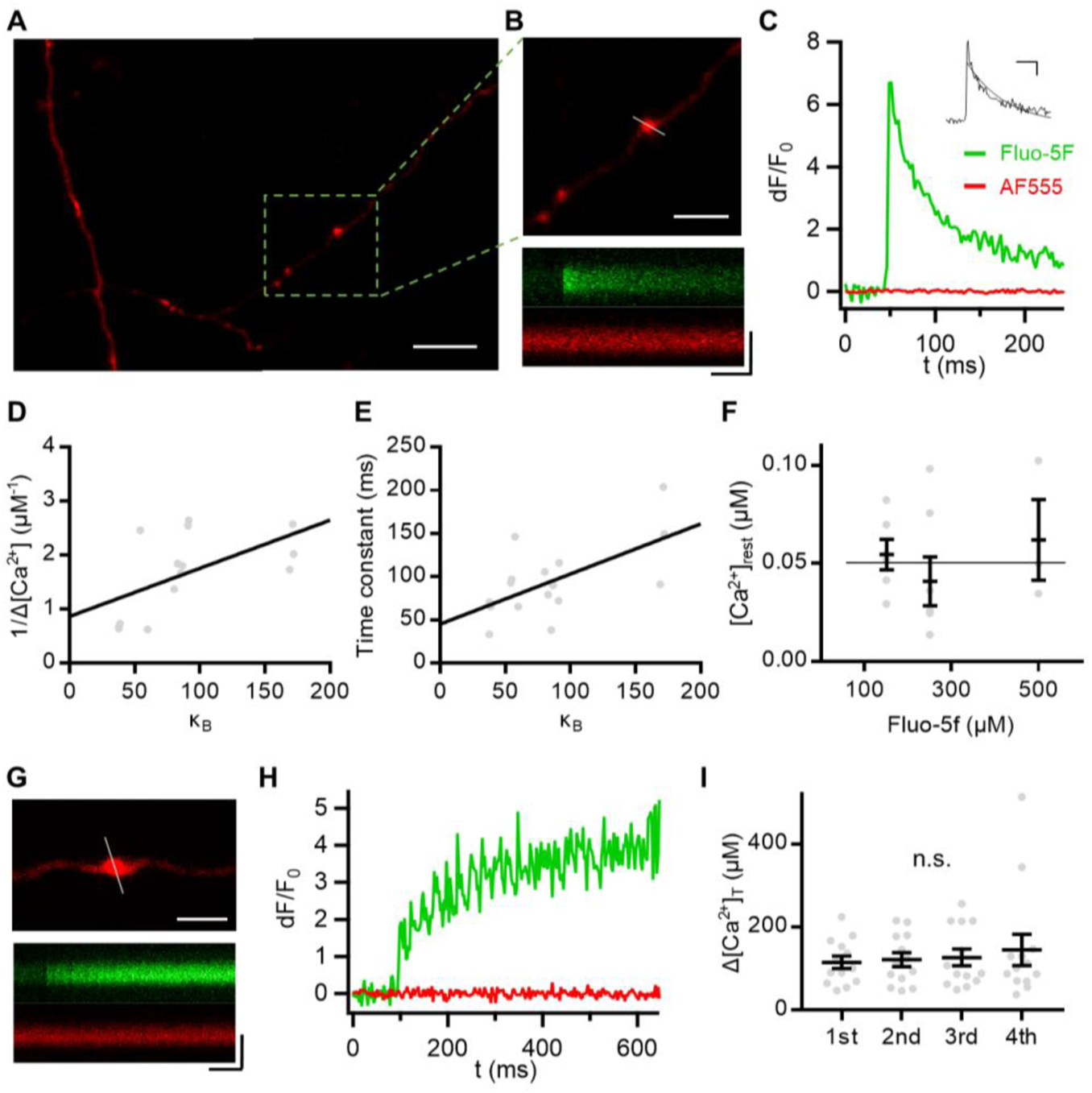
Presynaptic calcium kinetics at boutons of layer 2/3 pyramidal cells. (**A-C**) A mixture of Fluo-5F (150, 250 or 500 μM) and Alexa Flour 555 was loaded on their axonal boutons of L2/3-PCs through whole-cell patch techniques for dual excitation ratiometric measurements of the cytosolic [Ca^2+^] at axonal boutons. (**A**) The long-branched stretch of axon from L2/3 PCs visualized through maximum fluorescence intensity projection using confocal laser scanning of AF555 perfused via a patch clamp pipette. Scale bar, 10 μm. (**B**) *Top*, the selected bouton for recordings, among the three, is displayed with the corresponding scanning line. Scale bar, 5 μm. *Bottom*, Line-scan images for fluorescence of Fluo-5F and AF555. Scale bars, 50 ms and 2 μm. (**C**) Baseline-normalized fluorescence changes evoked by an AP. Fluorescence intensities were spatially averaged over the bouton shown in (*B*), background-subtracted and normalized to the baseline value (dF/F_0_). *Inset*, Ca^2+^ transient estimated from dF/F_0_. Scale bar, 50 ms and 100 nM. (**D, E**) The kinetic parameters of an AP-CaT were determined from a plot for the inverse of AP-evoked [Ca^2+^] increment (1/Δ[Ca^2+^]) *vs*. calcium binding ratio (κ_B_; Eq. 4 in *Material and Methods*). (**D**) Reciprocal plot of the AP-induced [Ca^2+^] changes (1/Δ[Ca^2+^]) vs. corresponding calcium binding constants (κ_B_) according to Eq. 2 (n = 14). A linear fit to the plot gave the amplitude of AP-CaT (A_Ca_ = 1.16 μM) from the y-intercept, representing the amplitude of free [Ca^2+^] increase in the absence exogenous Ca^2+^ buffer. The calcium binding ratio of endogenous static Ca^2+^ buffers (κ_S_) was estimated as 96 from the x-intercept of the linear regression based on Eq. 2 in *Material and Methods*. Linear regression line, P = 0.035. (**E**) Plot of [Ca^2+^] decay time constant versus κ_B_ (n = 16). The time constant for [Ca^2+^] decay from mono-exponential fit was plotted vs. κ_B_, and a linear regression based on Eq. 3 yielded the κ_S_ value of 77 from the x-intercept. The y-intercept was 43 ms, an estimate for a dye-independent value for Ca^2+^ decay time constant (τ_c_) (Eq. 3 in *Material and Methods*). Linear regression line, P = 0.011. (**F**) Resting [Ca^2+^] measured from basal fluorescence ratio using Eq. 1 in *Material and Methods* as a function of [Fluo-5F] in patch pipettes. The horizontal line indicates 50 nM. The mean [Ca^2+^]_rest_ at three different concentrations of Ca^2+^ indicator dye were not different and measured as 50 nM, suggesting that the dye did not influence [Ca^2+^]_rest_. (**G-I**) Strong PPD and the subsequent STF may be caused by Ca^2+^ channel inactivation and Ca^2+^-dependent facilitation (CDF) of calcium influx into presynaptic boutons, respectively. To test these possibilities, we measured presynaptic Ca^2+^ increments evoked by each of successive APs applied in a train at 40 Hz. (**G**) Fluorescence signals from a selected bouton with a scanning line. *Scale bars*, 3 μm (*top*), 100 ms and 2 μm (*bottom*). (**H**) Normalized fluorescence (dF/F_0_) responses to a train of APs at 40 Hz. (**I**) Total Ca^2+^ increments (Δ[Ca^2+^]_T_; free plus bound form) are plotted against the number of APs in the train (n = 13). Δ[Ca^2+^]_T_ were calculated based on Eq. 2 with κ_S_ set to 90. No significant change was observed for the estimates of Δ[Ca^2+^]_T_ at each of first 4 APs during 40 Hz train, indicating that contributions of Ca^2+^ channel inactivation and CDF to STP are unlikely. Mean ± S.E.M., One-way ANOVA; n.s. = not significant.

**Figure 3—figure supplement 1.**
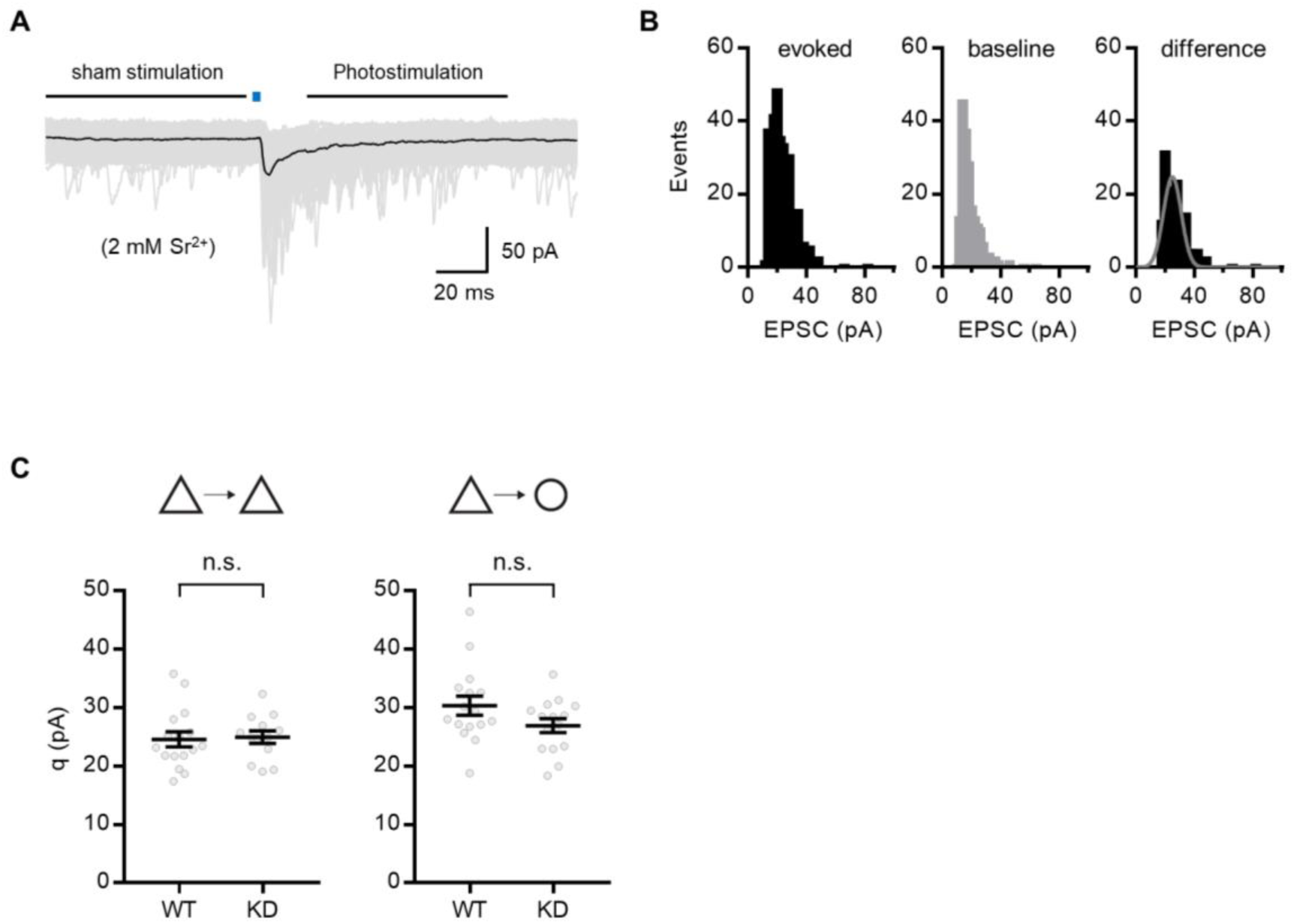
Quantal size estimated from Sr^2+^-induced asynchronous release. (**A**) Representative traces (100 pulses) from PC-PC synapses. Black bars indicate the detection window (100 ms) for release events in sham stimulation (*left bar*) and photostimulation (*right bar*). (**B**) Representative distribution of asynchronous EPSCs at PC-PC synapses. *Left*, after photostimulation. *Middle*, before stimulation. *Right*, difference between left and middle panels. The difference distribution was fitted with a Gaussian function (m = 24.93, σ = 6.09). (**C**) Comparisons of estimates for quantal size (q) between WT and Syt7 KD synapses. From left to right: PC-PC (n = 16, 13 for WT and KD, respectively), PC-FSIN (n = 16, 15). *Gray symbols*, individual data. All data are represented as mean ± S.E.M., unpaired t-test; n.s. = not significant.

**Figure 3—figure supplement 2.**
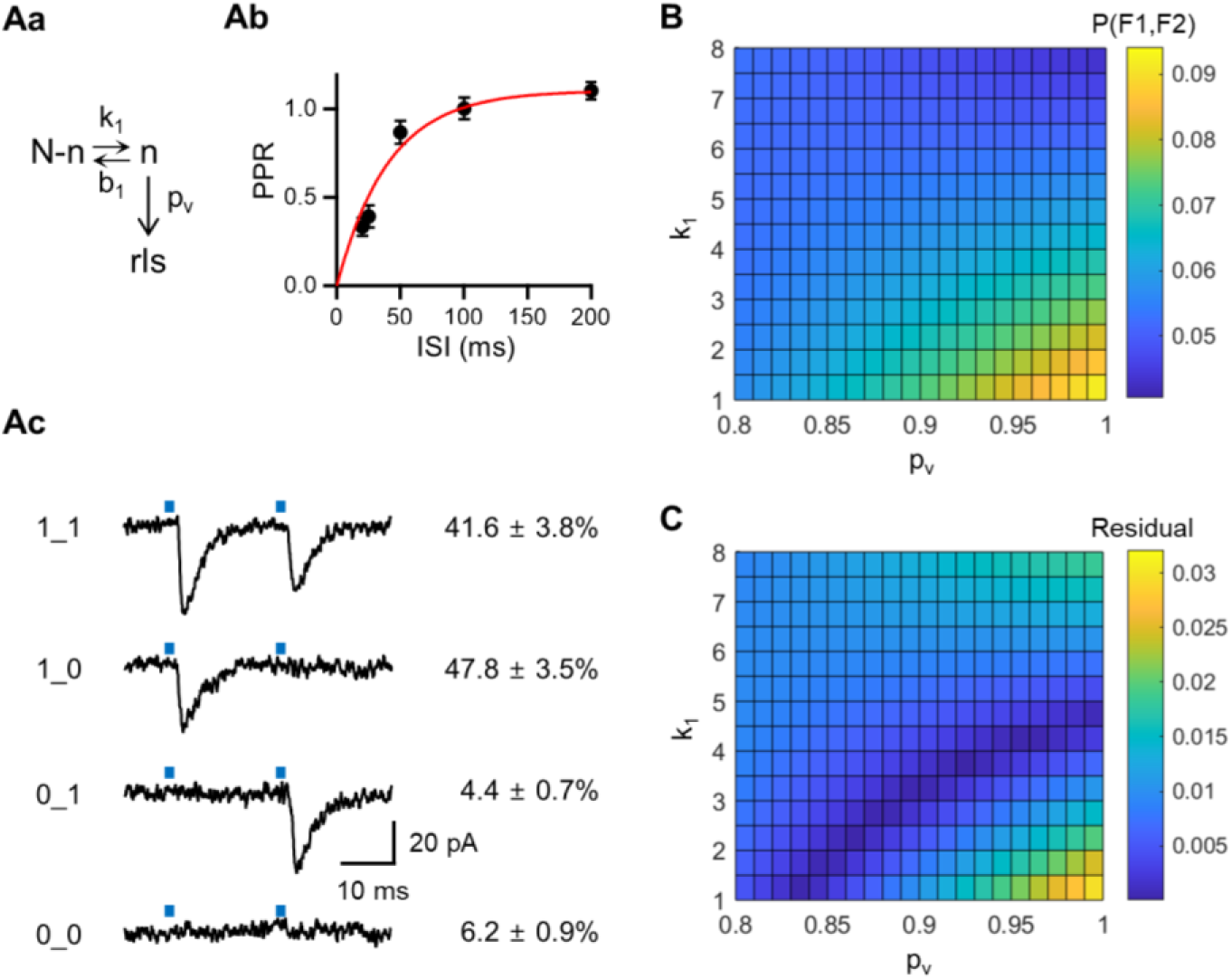
Analysis of double failure rate supports high p_v_ at excitatory synapses. (**Aa**) Schematic of simple refilling model. (**Ab**) An exponential fit to the PPR plot as a function of ISIs reproduced from Figure 1G. *Fitted function*, PPR = 1.1 [1 − exp(− t/43.5 ms)]. (**Ac**) Representative traces for four combinations of success and failure of 1st and 2nd EPSCs evoked by paired pulses with ISI of 20 ms at PC-PC synapses, showing successes in both pulses (1_1), failures in both pulses (0_0), and mixed responses (1_0; 0_1). On the right side of each trace, corresponding observation probability is shown in percentages (mean ± S.E.M. n = 57). (**B-C**) Plots of the calculated P(F1, F2) values (*B*) and their differences (*C*) from the observed value (6.2%) under the simple refilling model are displayed on the plane of k_1_ versus p_v_.

**Figure 3—figure supplement 3.**
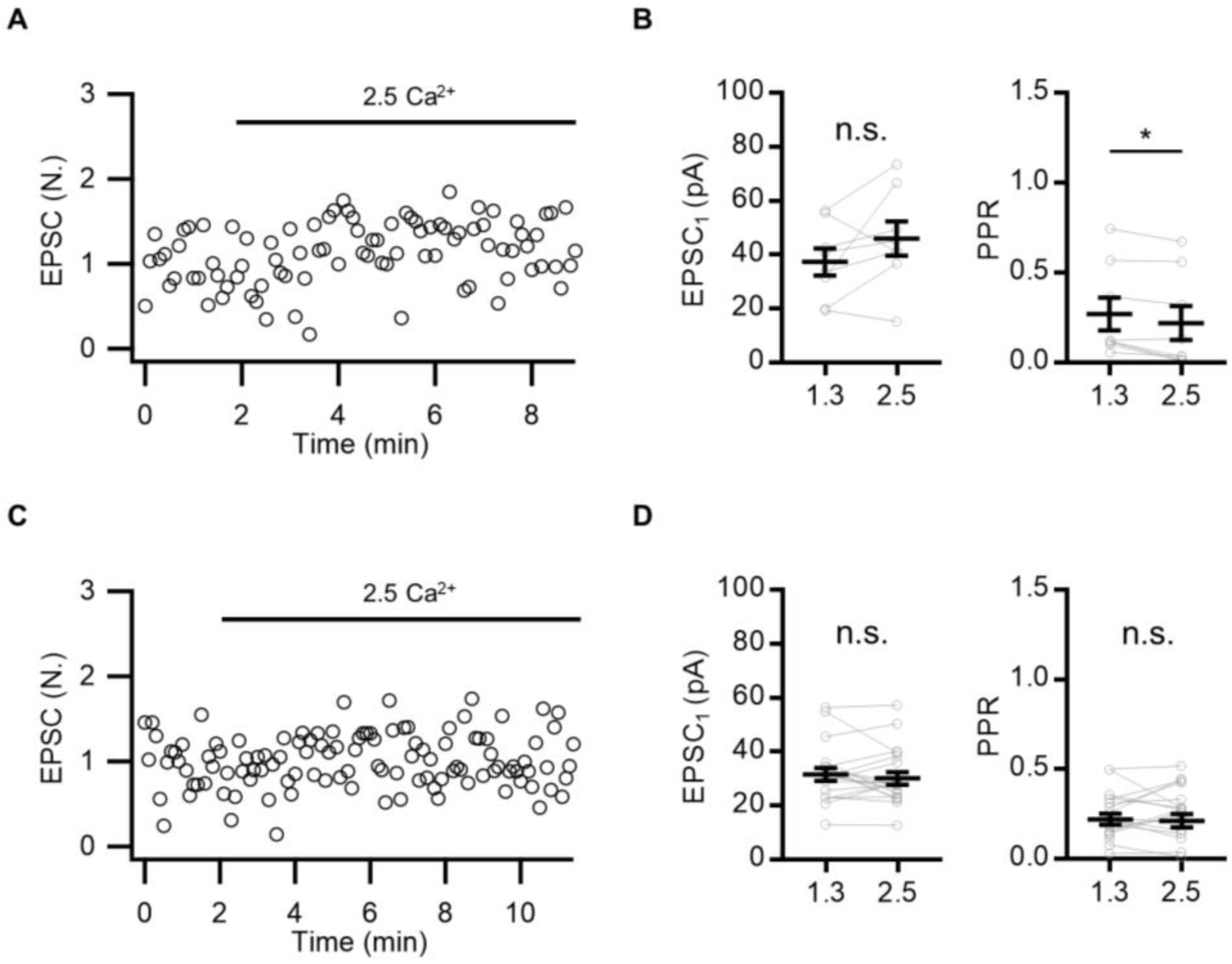
Minimal EPSC increase upon elevated [Ca²⁺]ₒ indicates saturated p_v_ at baseline. (**A**) Representative time courses of normalized amplitudes of the first EPSCs during paired-pulse photo-stimulation at 20 ms ISI under control conditions (1.3 mM [Ca²⁺]ₒ) and elevated extracellular [Ca²⁺] (2.5 mM; n = 8). (**B**) Mean values for the first EPSC (*left*) and PPR (*right*) under the two conditions. (**C-D**) Experiments of A-B were repeated in the slices that were pre-incubated in aCSF containing 50 μM EGTA-AM (n = 20). Note that the modest extracellular [Ca^2+^]-dependence of the first EPSC and PPR observed in A-B was abolished by 50 μM EGTA-AM pre-incubation. *Gray symbols*, individual data. All statistical data are represented as mean ± S.E.M.; n.s. = not significant; *, P<0.05; paired t-test.

**Figure 5—figure supplement 1.**
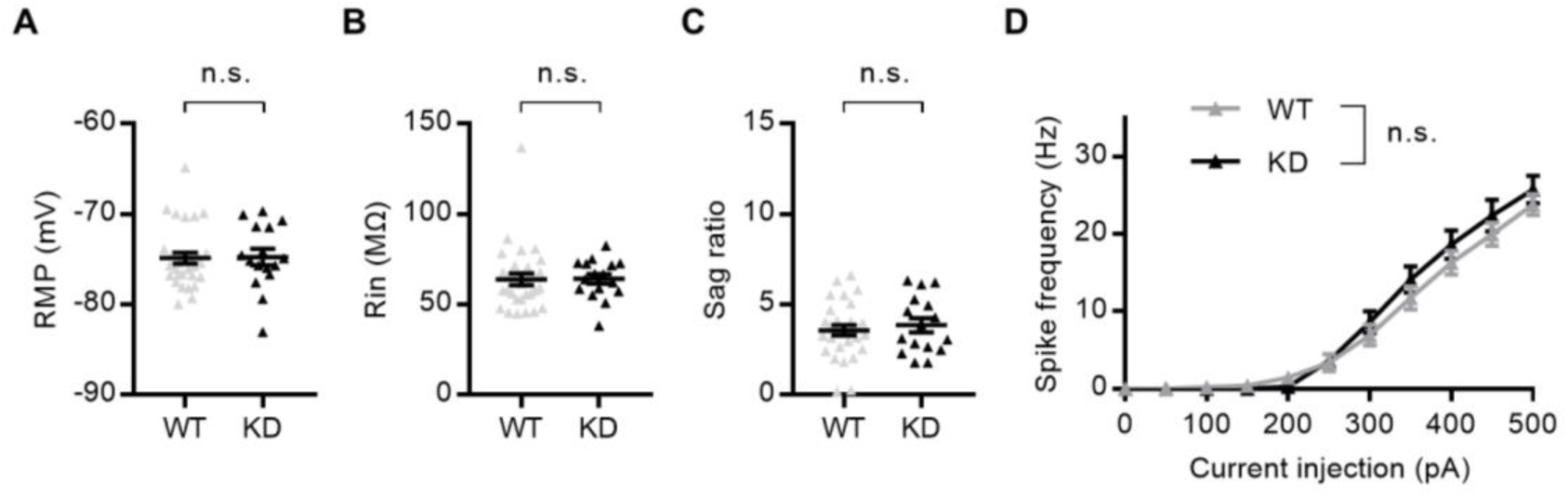
Little effects of Syt7 KD on intrinsic properties of neurons. Whole-cell recordings were performed from the fluorescently labeled pyramidal neurons in the prelimbic L2/3 of WT (n = 30) or Syt7 KD (n = 16) rats. (**A**) Resting membrane potential. (**B**) Input resistance. (**C**) Sag ratio. *Filled triangles*, individual data. (**D**) Current-spike frequency relationship in WT and KD cells. All statistical data are represented as mean ± S.E.M., unpaired t-test or two-way ANOVA; n.s. = not significant.

**Figure 5—figure supplement 2.**
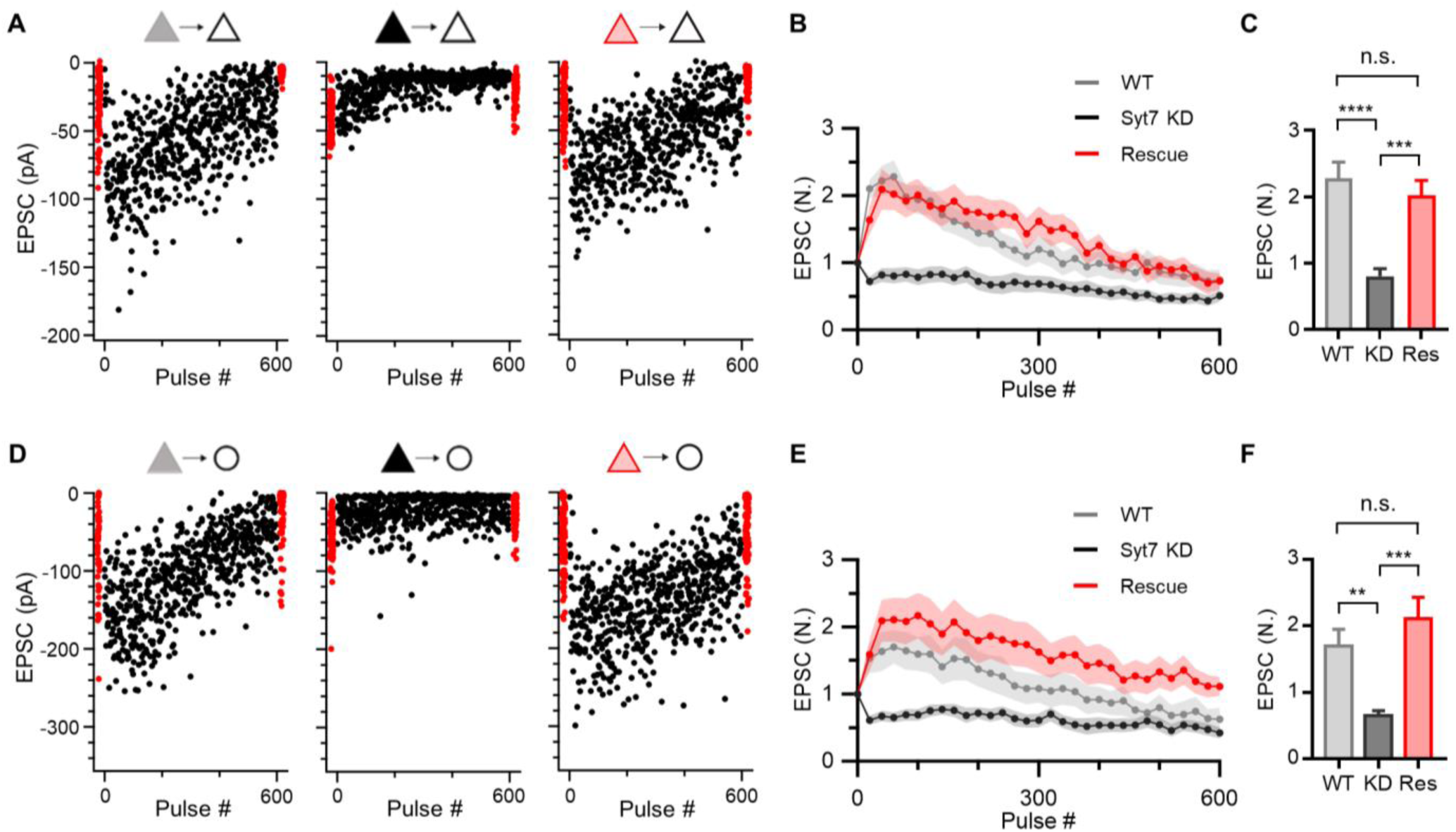
Presynaptic expression of Syt7 rescues facilitation at L2/3 excitatory synapses. (**A, D**) Representative amplitudes of EPSC trains evoked by 600-pulse 10 Hz trains (flanked by 2 baseline periods comprised of 40 pulses each) at PC-PC (*A*) and PC-FSIN (*D*) synapses, in which presynaptic WT (*left, gray triangles*) pyramidal cells and those transfected with plasmid encoding shRNA against Syt7 (sh-Syt7, KD, *middle, black triangles*) or sh-Syt7 plus Syt7 insensitive to shRNA (rescue, *right, red triangles*). (**B, E**) STP of EPSCs evoked by 600-pulse trains at 10 Hz at L2/3 PC-PC (*B*; n = 9, 10, 7; WT, Syt7 KD, Rescue, respectively) and PC-FSIN (*E*; n = 12, 10, 8) synapses. Each data point represents an average from a moving window of 20 pulses, normalized to the baseline EPSC. (**C, F**) Mean values of 41-60^th^ pulses from *B* and *E*. Mean ± S.E.M.; n.s. = not significant; **, P<0.01; ***, P<0.001; ****, P< 0.0001; one-way ANOVA with Tukey’s post-hoc test.

**Figure 8—figure supplement 1.**
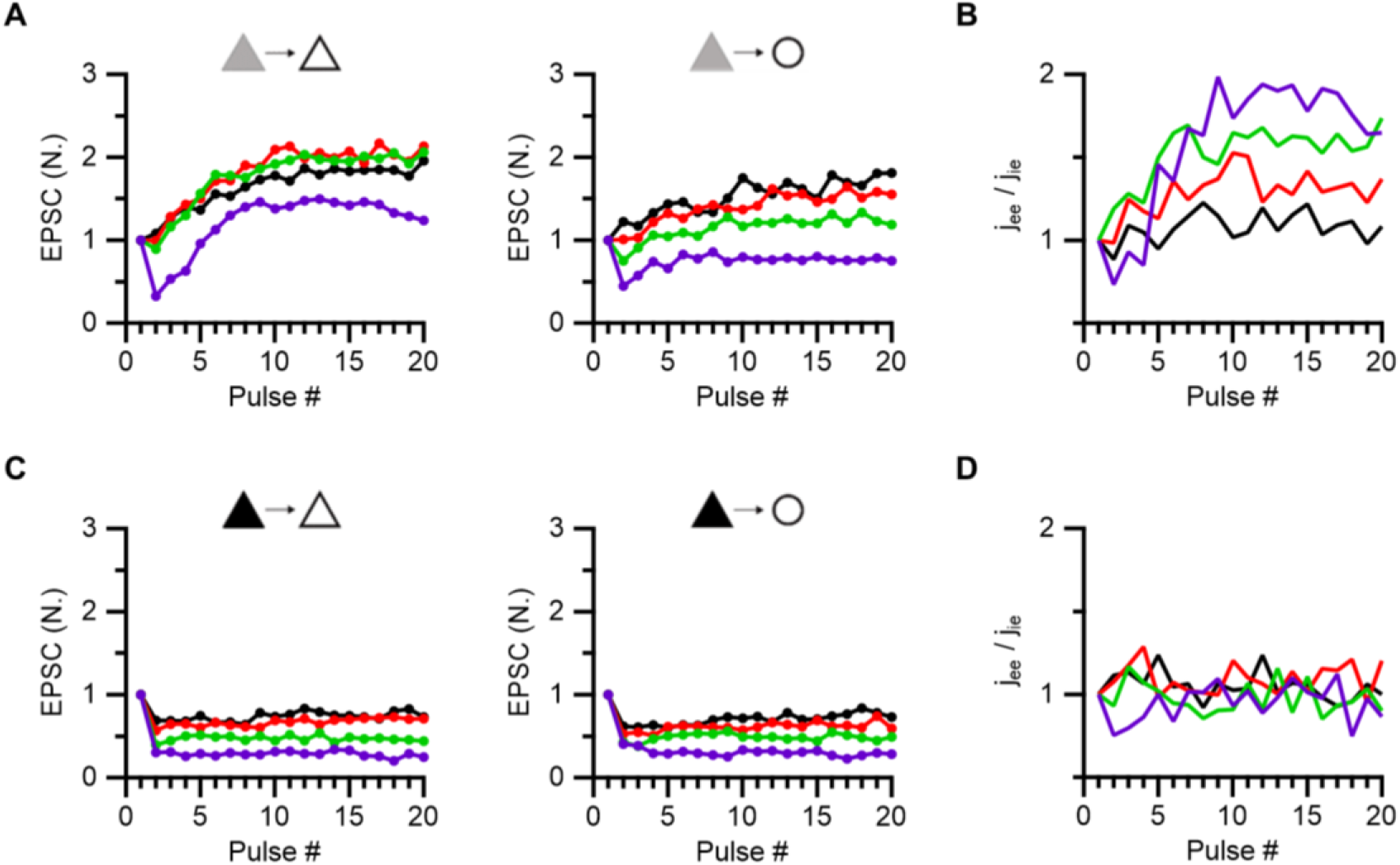
Activity-dependent increase in the synaptic weight ratio at PC-PC to at PC-FSIN connections (J_ee_/J_ie_) is abolished by Syt7 KD. (**A, C**) Average amplitudes of baseline-normalized EPSCs during 20-pulse trains at frequencies from 5 to 40 Hz at PC-PC (*left*) and PC-FSIN (*right*) synapses in WT (*A*) and Syt7 KD (*C*). Data are reproduced from Figure 1 (*A*) and Figure 5 (*C*). (**B, D**) Relative synaptic weights of excitatory input to PCs over that onto FSINs (*Jee/Jie*) in WT (*B*) and Syt7 KD (*D*). Note that excitatory influence (*Jee/Jie*) to the recurrent network increases as the firing number and frequency increase in WT but not in Syt7 KD.

**Figure 8—figure supplement 2.**
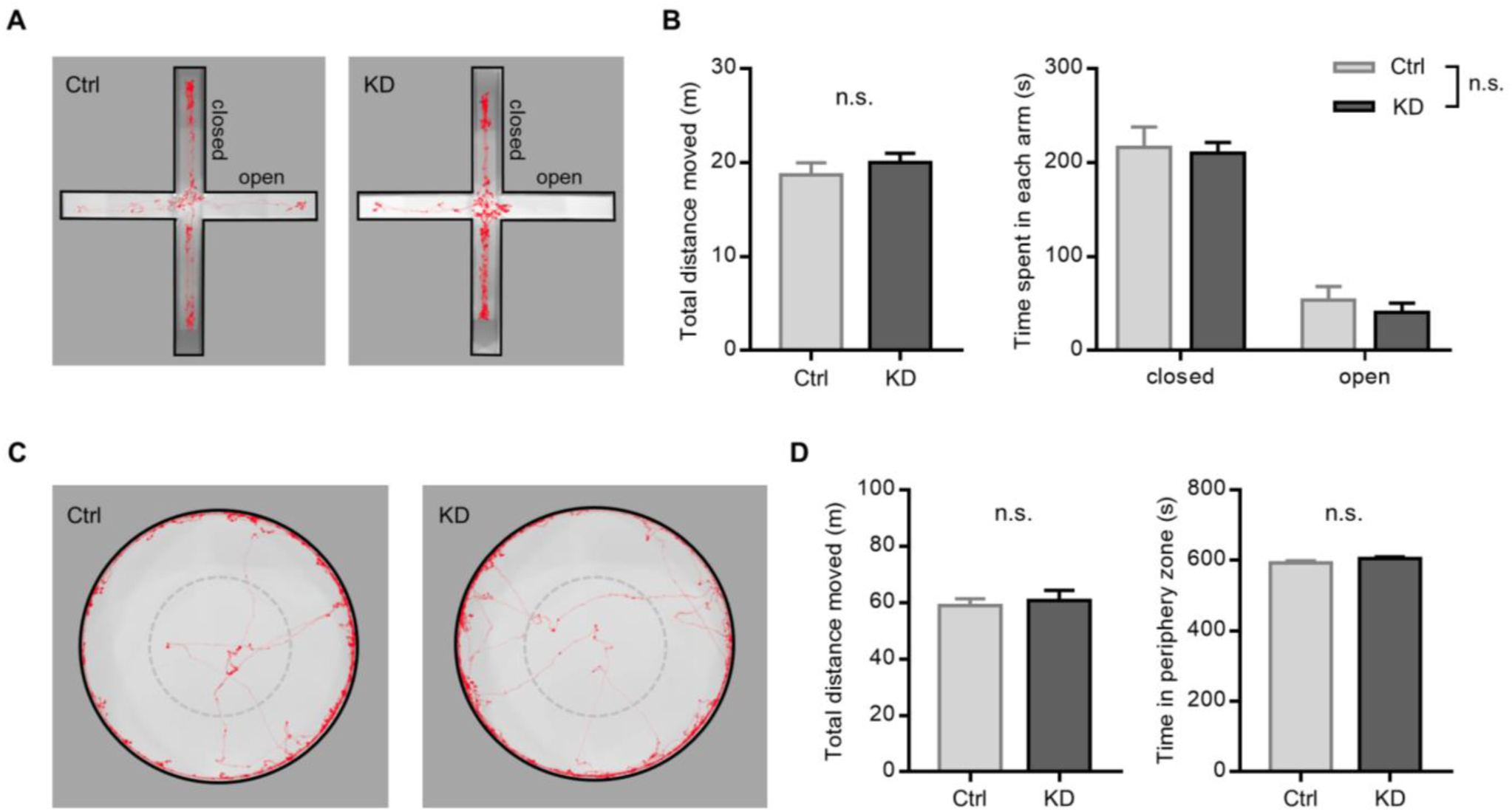
Locomotor activity and anxiety levels in Syt7 KD rats using OFT and EPM. (**A, C**) Representative tracking trace of the elevated plus maze (*A*) and open field test (*C*). (**B, D**) Mean values for total distance moved (*left*) and time spent (*right*) in each arm (*B*) or in periphery zone (*D*) in control (n = 13) or Syt7 KD rats (n = 9). Mean ± S.E.M.; Unpaired t-test or two-way ANOVA; n.s. = not significant.

